# Collective cell migration without proliferation: density determines cell velocity and wave velocity

**DOI:** 10.1101/232462

**Authors:** Sham Tlili, Estelle Gauquelin, Brigitte Li, Olivier Cardoso, Benoît Ladoux, Hélène Delanoë-Ayari, François Graner

## Abstract

Collective cell migration contributes to morphogenesis, wound healing or tumor metastasis. Culturing epithelial monolayers on a substrate enables to quantify such tissue migration. By using narrow strips, we stabilise the front shape; by inhibiting cell division, we limit density increase and favor steady migration; by using long strips, we observe a confined cell monolayer migrating over days. A coherent collective movement propagates over millimeters; cells spread and density decreases from the monolayer bulk toward the front. Cell velocity (∼micrometer per minute) increases linearly with cell radius, and does not depend explicitly on the distance to the front. Over ten periods of backwards propagating velocity waves, with wavelength ∼millimeter, are detected with a signal-to-noise ratio enabling for quantitative spatio-temporal analysis. Their velocity (∼ten micrometers per minute) is ten times the cell velocity; it increases linearly with the cell radius. Their period (∼two hours) is spatially homogeneous, and increases with the front density. When we inhibit the formation of lamellipodia, cell velocity drops while waves either disappear, or have a smaller amplitude and slower period. Our phenomenological model assumes that both cell and wave velocities are related with the activity of lamellipodia, and that the local stretching in the monolayer bulk modulates traction stresses. We find that parameter values close to the instability limit where waves appear yield qualitative and quantitative predictions compatible with experiments, including the facts that: waves propagate backwards; wave velocity increases with cell radius; lamellipodia inhibition attenuates, slows down or even suppresses the waves. Together, our experiments and modelling evidence the importance of lamellipodia in collective cell migration and waves.

## Introduction

Collective migration of cells connected by cell-cell adhesion occurs across large time scales and length scales in numerous biological processes like embryogenesis (notably gastrulation), wound healing, regeneration or tumor metastasis (1–4). To study such mechanical behaviour of tissues, *in vitro* reconstructed assemblies of cohesive cells are useful experimental model systems (5, 6) where each individual cell retains its normal physiological behavior: it can grow, divide, die, migrate. In two-dimensional (2D) monolayers, cells interact with each other biochemically and mechanically, for instance through adhesion, and have a richer migration behaviour than single cells. It is possible to constrain geometrically and reproducibly control their collective migration. Patterned substrate of adhesive strips enable to investigate the tissue global response to active processes like cell migration (5, 7) or cell division (8), and quantitatively test the impact of drugs such as blebbistatin (9). Madin-Darby canine kidney (MDCK) cell monolayers enable comparisons of experiments, simulations and theories (10–15); 2D images are easier to obtain and analyze, to extract physical quantities such as cell velocity, density, shape and deformation (12, 16).

When monolayers are grown on a substrate, it acts as a source of external friction on cells (5, 7, 11, 17). If it is deformable (made of soft gel or covered with pillars), it acts as a mechanical sensor for traction force microscopy to quantify forces exerted by cells on the substrate, which are the opposite of forces exerted by the substrate on the cells (18–20). Beside these external forces, mechanical stresses within the monolayer arise from cell-level processes which include: cell volume change (21) and division (8); competition between the adhesion to the substrate, the intercellular adhesion and the cell contractility (22); cryptic lamellipodia extending from one cell below its neighbours (23).

The emergence of large-scale polarized movements within epithelial cell monolayers largely depends on mechanical factors and external geometrical constraints (7, 13, 16, 24). Loza et al. (using human breast epithelial cells) showed that cell density and contractility control transitions in collective shape, and could predict *in vivo* collective migration in a developing fruit fly epithelium (25). Microfluidic channel experiments have shown that the flow velocity of the front can be decomposed into a constant term of directed cell migration superimposed with a diffusion-like contribution that increases with density gradient (26). In the context of a cell monolayer collectively spreading and invading a free space, highly motile leader cells can appear (27) and locally guide small organized cohorts of cells (10). The cell velocity decreases with the distance to the moving front (11) while both the cell density and the stress increase with the distance to the moving front (5). Bulk cellular motions also display large-scale coordinated movements of cell clusters that can be seen by the emergence of typically 200 *μm* correlation length for the velocity field and large-scale polarization (9, 28).

Serra-Picamal *et al.*, by confining cells on a strip then releasing the confinement, observed two periods of a mechanical wave, propagating backwards from each front, made visible by oscillations of the cell velocity and its gradient, and suggesting how stress mediates collective motion (11). Mechanical force propagation has been reported during the collision of two epithelial cell layers to explain the formation of tissue boundaries (29). Similar wound healing experiments displayed a wave of coordinated migration, in which clusters of coordinated moving cells were formed away from the wound and disintegrated near the advancing front; this wave could be amplified by the hepatocyte growth factor / scatter factor (30). Confluent epithelial cells confined within circular domains exhibit collective low-frequency radial displacement modes as well as stochastic global rotation reversals (31, 32). While oscillations at smaller scales are common in embryogenesis (cell size and minute period (33–36)) or myxobacteria swarms (a few cell sizes, 1 to 100 minutes period (37)), here in confluent monolayers the oscillation scale is that of a tissue size and of hours, reminiscent of somitogenesis (for review of models, see Ref. (38)).

Even though the appearance of cell coordination and waves in collective migration experiments is crucial to understand important biological functions, it remains poorly understood. Migration and division contributions to the front velocity are entangled. Moreover, cell number is constantly increasing due to cell division, which leads to jamming and slowing of the migration. This usually limits the experiment duration to a few hours. To improve our understanding, distinguish between the models, and constrain their parameters, varied and controlled experimental data are required.

Here, by inhibiting cell division, we rather see a decrease in cell density due to migration. We observe a steady collective migration over a day or more, without reaching jamming densities. The strip length is adapted to such long-distance migration. Strips are narrow to prevent front shape instabilities, and the cell flow is essentially uni-dimensional (Fig. 1, Fig. S1 and Movies S1-6 in the Supporting Material). Backwards propagating velocity waves are detected. We first average out the velocity field on a timescale larger than a wave period, to characterize the mean cell velocity profile in the monolayer bulk: it depends explicitly only on the cell density (it increases linearly with the mean effective cell radius), irrespectively of the distance to the migrating front. We then quantify the waves with an unprecedented signal-to-noise ratio; their velocity and pulsation decrease with cell density. Inhibiting lamellipodia formation drastically decreases the monolayer average velocity; it decreases the pulsation and amplitude of the waves or even leads to their suppression. Finally, a phenomenological model of collective migration suggests how to interpret these experimental observations.

**Figure 1:**
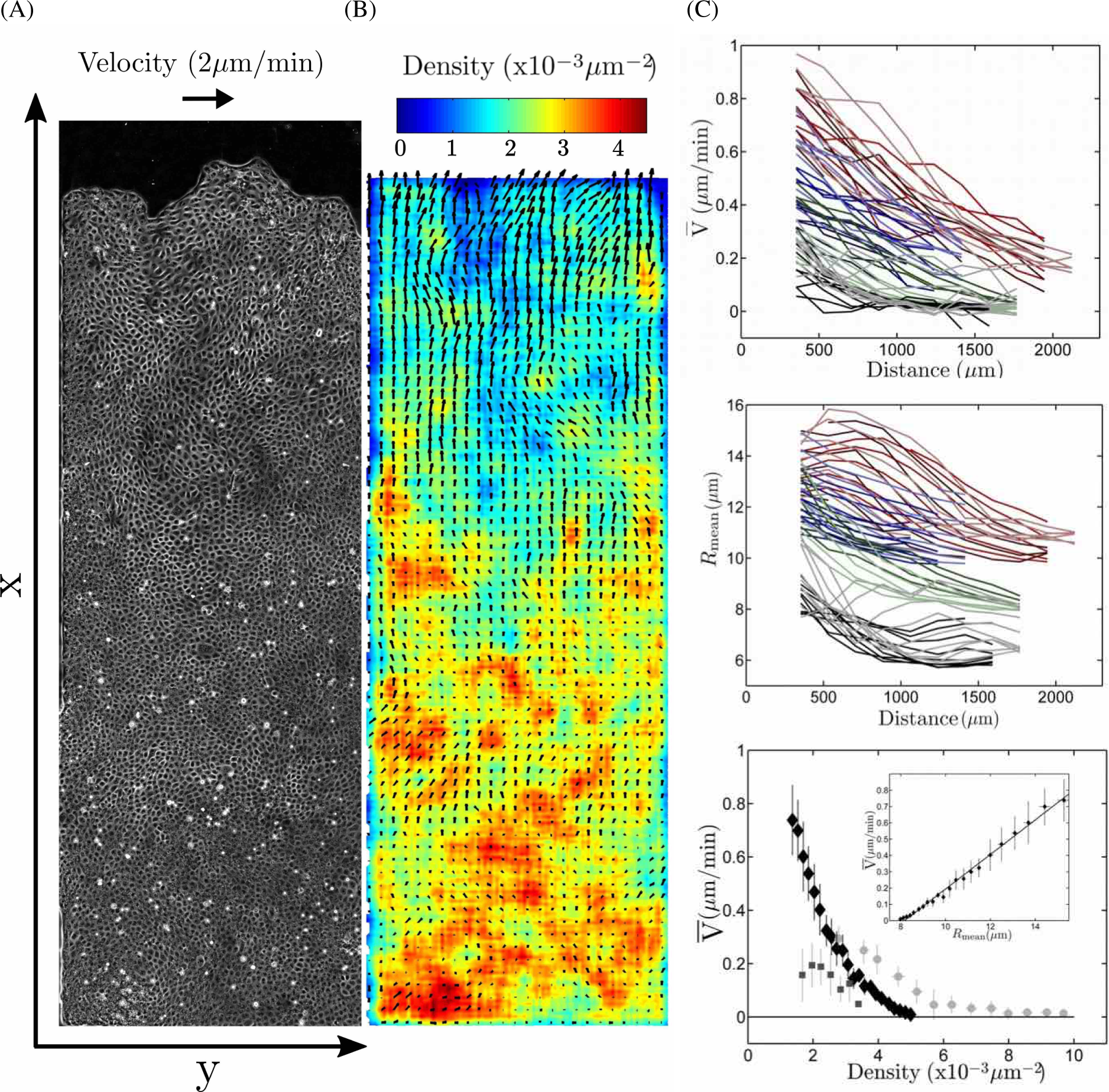
Cell migration. (A) A monolayer of MDCK cells, initially confined, is released. It expands (Movies S1-3) along the adhesive strip towards empty space (direction of increasing *x*). Mitomycin C is added to inhibit divisions. Phase-contrast image of cell contours, taken at *t* = 11 h 30 min (i.e. after ∼16 h 30 min of migration). Strip total length 4 mm (most of it is visible here), width 1 mm. (B) Corresponding 2D fields of cell velocity and density. Scale arrow: 2 *μ*m/min. (C) Large scale profiles. Large-scale average of cell velocity 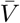 (top) and mean effective cell radius 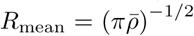 (middle), plotted vs distance *X* to migrating front. Each color marks a different batch, with red, blue, green in order of increasing initial density. For a given color (i.e. batch), each shade marks a different strip. For a given shade (i.e. strip), each data point is the average in a box of 176 *μ*m × 180 min. Grey levels are the same, for control experiments without mitomycin C. Bottom: Same data, with mitomycin, binned and plotted as 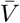 vs cell density 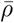 (black diamonds) or vs *R*_mean_ (inset), with a linear fit 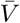 = 0.106 *R*_mean_ – 0.864 (R=0.9931); *N* = 8 strips. Light grey circles: control experiments without mitomycin C; *N* = 3 strips. Grey squares: experiment with CK666 to inhibit lamellipodia; *N* = 5 strips. Horizontal and vertical bars: standard deviation (SD) within each bin.

## Materials and methods

### Micropattern printing

The micropattern of fibronectin is printed according to the following standard soft lithography technique, robust to small changes in the procedure. Patterned PDMS stamps prepared from silanized wafers are incubated for 45 min at 37°C or 60 min at room temperature with a solution of 50 *μ*g/mL of fibronectin (Sigma) and 25 *μ*g/mL of Cy3 conjugated fibronectin. A thin layer of PDMS (10 % of reticulating agent) is spin coated on a 35 mm plastic Petri dish and cured for 2 h at 80°C or overnight at 65°C. The Petri dish is exposed to UV for approximately 20 min in order to activate the PDMS surface. After incubation, stamps are dried and pressed on the UV activated PDMS surface in order to transfer fibronectin. A 2 % Pluronic F-127 (Sigma) solution is added to the Petri dish to chemically block the regions outside of the fibronectin pattern for 1 hour at room temperature. The Pluronic solution is removed after 1 hour and the Petri dish is rinsed 3 to 6 times with a PBS solution.

### Cell culture

We use the same MDCK strain as in Ref. (9). A stable cell line was created using Histone GFP (39) using the DreamFect Gold Transfection Reagent, Oz Biosciences.

A batch has 3 to 6 strips, with the same initial MDCK cell density, lengths up to 4 mm. Strip widths range from 200 *μ*m to 1 mm (at least equal to the typical 200 *μ*m correlation length for the velocity field (9, 28)) and do not affect the results presented here. Different batches correspond to different initial cell densities, tested at least twice each. Suspended cells are deposited and allowed to attach for one to a few hours. Non-attached cells are rinsed, while attached cells grow and divide until full confluence. The confining PDMS block is removed. Some cells might detach, so the monolayer is rinsed again and left for a few hours. The monolayer starts to migrate along the whole accessible strip, expanding towards the empty surface where cells adhere to fibronectin, and not towards outside regions chemically blocked using Pluronic.

In order to decrease the division rate, 8 *μ*L of a 0.5 mg/mL mitomycin C solution is added to 1 mL cell culture medium, and cells are incubated at 37°C for 1 h (31, 40). They are then abundantly rinsed with fresh 37°C medium to prevent the toxicity effects reported for 12 h exposure to mitomycin (28). After 3 h the division rate is less than a fifth of the initial one (Fig. S2 in the Supporting Material), and the rate of extrusions also strongly decreases. Control experiments in standard conditions, with proliferating cells (no mitomycin C added), are performed with initial cell density 5 10^−3^ *μ*m^−2^.

To test the role of lamellipodia, we prepare a 100 *μ*M solution in DMSO of CK666, namely 2-Fluoro-N-[2-(2-methyl-1H-indol-3-yl)ethyl]benzamide, a selective inhibitor of actin assembly mediated by actin-related protein Arp2/3 (IC50 = 17 *μ*M) (41–43). Aliquots are stored at −20°C and used within two weeks of preparation. The solution is added to the cells after ∼ 1 day of migration and is not rinsed. Lamellipodia (both cryptic and front ones) are no longer detectable (Movie S7 in the Supporting Material).

### Imaging

Two hours after having added the mitomycin, we take the first image of the movie and define it as *t* = 0. Live imaging of monolayers is performed in the Nikon BioStation IM, a compact cell incubator and monitoring system, with an air objective (CFI Plan Fluor 10X, Nikon). Phase-contrast and fluorescent imaging are used to observe respectively cell contours and cell nuclei. The interframe time interval is 5 min for 1 mm wide strips and 6 min for 200 *μ*m wide strips.

Dead or extruded cells appear as bright spots which can be removed by manual image intensity thresholding. The contrast is adjusted separately on each color channel, and a blur with 2 pixel radius removes sharp intensity fluctuations. To obtain the whole view of the confined monolayer, up to 6 (for two 200 *μ*m wide strips) or 20 (for 1 mm wide strips) microscope fields of view are merged using the Grid/Collection Stitching Plugin (44) implemented in ImageJ. We use the “unknown position” option for the first time frame to calculate automatically the overlap between images, which we use for all frames since images are stable.

### Velocity and density

We measure the two-dimensional velocity field 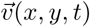 (Fig. 1B and Fig. S1A,C) using Particle image velocimetry (45). We use the open source toolbox MATPIV *matpiv.m* (46) of Matlab (The MathWorks, Inc., Natick, Massachusetts, United States), with the “singlepass” option, square box of side 32 pixels (20 *μ*m) for 200 *μ*m wide strips and 128 pixels (80 *μ*m) for 1 mm wide strips, and box overlap is 50 % or 75 % for both widths.

The Particle image velocimetry method, option “single”, interrogation box size of 128 pixels, yields qualitatively identical results, and is quantitatively around 10% larger, when compared either with “multin” option, windowsize-vector [128 128;64 64] or with Kanade-Lucas-Tomasi (KLT) feature matching algorithm, pyramid parameter 2, successive interrogation box sizes of 128 and 64 pixels.

We do not detect any statistically significant dependence of 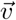 with *y*, even near the lateral sides of the strip. The y component of 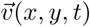 is lower at higher positions *x* where the average velocity is higher (Fig. S1C), indicating a more directed movement; we do not consider this component in what follows. The component of 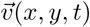 along the *x* axis, i.e. along the long axis of the strip, averaged over *y*, is the one-dimensional velocity field *V*(*x, t*), which we study here.

To plot the space-time diagram or “kymograph” of *V*(*x, t*), we remove noise using a Gaussian blur of standard deviation 15 min and 30 *μ*m (and a sliding window which is three times larger). We then separate scales, and decompose *V* into 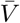 and V – 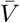, using a Gaussian filter of standard deviation 50 min and 100 *μ*m (again, with a sliding window which is three times larger). For large scale profiles 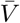, discrete measurements used for graphs are performed with an average on time t over 180 min and on space over 176 *μ*m wide bins in the distance *X* to the moving front. Only significant data points are plotted, i.e. points with enough pixels in the 176 *μ*m × 180 min box (at least 150 pixels, out of a maximum of 612) and where the signal value is larger than its SD. In particular, we entirely exclude the first box, where statistics are noisy due to the front. For the velocity gradient, a finite difference gradient is used and the resulting very small scale noise is removed with a 3-pixels wide linear filter.

Similarly, by using histones (Fig. S1B), we identify cell nuclei. Our measurements are based on local maxima detection, independently of the maximal intensity value, and thus are not sensitive to possible variations in the intensity of the GFP signal. The nuclei density can vary by a factor 10 within a given image. We first use a low blur radius, optimised for the highest nucleus density on the image. It yields a good detection for high density regions using *FastPeakFind.m* but gives false positives (more than one local maxima per nucleus) for low density regions. When the distance between two maxima is smaller than a critical value (equal to a third of the the local average distance between nuclei), we remove the less intense one. According to manual checks on high, middle and low density regions, the precision is better than 5 %.

We have checked that tracking the cell nuclei yields more fluctuations than PIV for 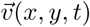 measurements, due to intracellular movements of nuclei (Movies S3-6), but yields same results as PIV for *V*(*x, t*). We use this cell nuclei detection to plot the cell density *ρ*(*x, t*), using boxes (*x, y*) of 20 *μ*m × 20 *μ*m, then an average over *y*. Using the same filters as for *V*, we remove noise, decompose *ρ* into 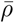 and *ρ* – 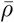, and define discrete measurements of *ρ* versus *X*.

### Wavelets

Kymographs are analysed using wavelet transform profilometry (WTP), a method inspired by 3D-fringe projection profilometry (47,48). It involves a one-dimensional continuous wavelet transform with a phase estimation approach. It is reliable, easy to implement, and robust to noise. We choose a Morlet wavelet and the wavelet transform is computed using a FFT algorithm (which is equivalent to an analytic Morlet Wavelet) with a Matlab script (49).

For each kymograph line (i.e. for each fixed position *x_i_*), the signal wavelet transform is computed at various time scales, in an observation window, 80 to 400 min, chosen in order to cover the full range of characteristic times of the observed oscillations (we checked that this choice does not affect the results presented here). The wavelet transform returns a matrix of complex coefficients *A*(*x_i_*, *t, s*), defined as continuous wavelet coefficients where *s* represents the test times scales. Each coefficient provides a local measurement of the similarity between the signal and the wavelet at a scale *s*. For each point (*x_i_*, *t_j_*) a given line *i* in the kymograph, only the coefficient *A_m_* (*x_i_*, *t_j_*) having the largest modulus with respect to the scale *s* is kept.

The argument of *A_m_*(*x_i_*, *t_j_*) provides the wrapped fringe phase *ϕ_w_*(*x_i_*, *t_j_*). The phase *ϕ_w_*(*x_i_*, *t_j_*) is unwrapped along time, and the local pulsation *ω* is deduced by differentiation with respect to time *t*. Independently, the phase *μ*(*x_i_*, *t_j_*) is unwrapped along space, and the local wave number *k* is deduced by differentiation with respect to space *x*.

## Results

### Large scale profiles of velocity and density

By averaging over the direction *y* perpendicular to the stripes, we determine the one-dimensional cell velocity field *V*(*x, t*) and cell density field *ρ* (*x, t*). We first investigate their overall profiles 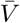, 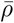, obtained by large scale sliding average. Due to the spreading of cells, variations of velocity and density are visible (Fig. 1). Far from the front (bottom of Fig. 1B), the density is still close to its initial value and the velocity is still zero, while close to the front (top of Fig. 1B) the density has decreased and the velocity increased. Note that far from the front, velocities are occasionally negative. We introduce *R* = (*πρ*)^−1/2^, then 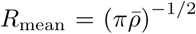, interpreted as a mean effective cell radius. Its typical range of variation is 8 to 15 *μ*m; for comparison, note that 4.8 *μ*m corresponds to the nuclei being almost close packed, while 17 *μ*m is the radius of a front cell at the limit of detaching from the monolayer.

With respect to the distance *X* to the moving front, different experimental batches display 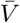(*X*) profiles which are qualitatively similar and quantitatively different (Fig. 1C, top), and *R*_mean_(*X*) profiles too (Fig. 1C, middle). When eliminating the space variable *X*, points coming from different batches fall on the same curve: 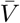 has a strong, negative correlation with 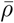 (13) (Fig. 1C, bottom; Fig. S3A in the Supporting Material), decreasing from 0.8 *μ*m/min to 0 for 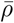 ranging from 1.8 to 5 10^−3^ *μ*m^−2^. In fact, 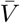 increases linearly with R_mean_ (Fig. 1C, bottom, inset). This relation does not depend on the distance to the front, and it is unaffected when the sliding window size in time is doubled.

We also performed experiments under standard conditions (i.e. without mitomycin C, Fig. S4 in the Supporting Material). While the overall relation between 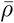 and 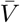 seems qualitatively unaffected, dividing cells have a significantly larger 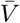 at given 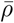, and a larger arrest density: 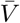 decreases from 0.3 *μ*m/min to 0 for 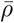 ranging from 3 to 10 10^−3^ *μ*m^−2^ (Fig. 1C, bottom; Fig. S3A).

### Propagating waves

We now turn to smaller scale variations. The cell velocity *V*(*x, t*) displays waves: cells slow down and accelerate while waves propagate from the front backwards in the –*x* direction (Movies S1-6). In the moving frame of average cell velocity, these waves would appear as a periodic velocity reversal. They are visible quantitatively onthe kymograph (Fig. 2A, Fig. S5A in the Supporting Material), and even more clearly on the velocity small scale variations 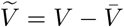 (Fig. S6A in the Supporting Material) as well as on the velocity gradient (Fig. 2B). The waves are reproducibly observed near the front, with a good signal-to-noise ratio over more than ten periods for a whole observation duration of *μ* 20 h. We do not detect any particular effect of the strip width on waves (compare Figs. S4B, S5A).

**Figure 2:**
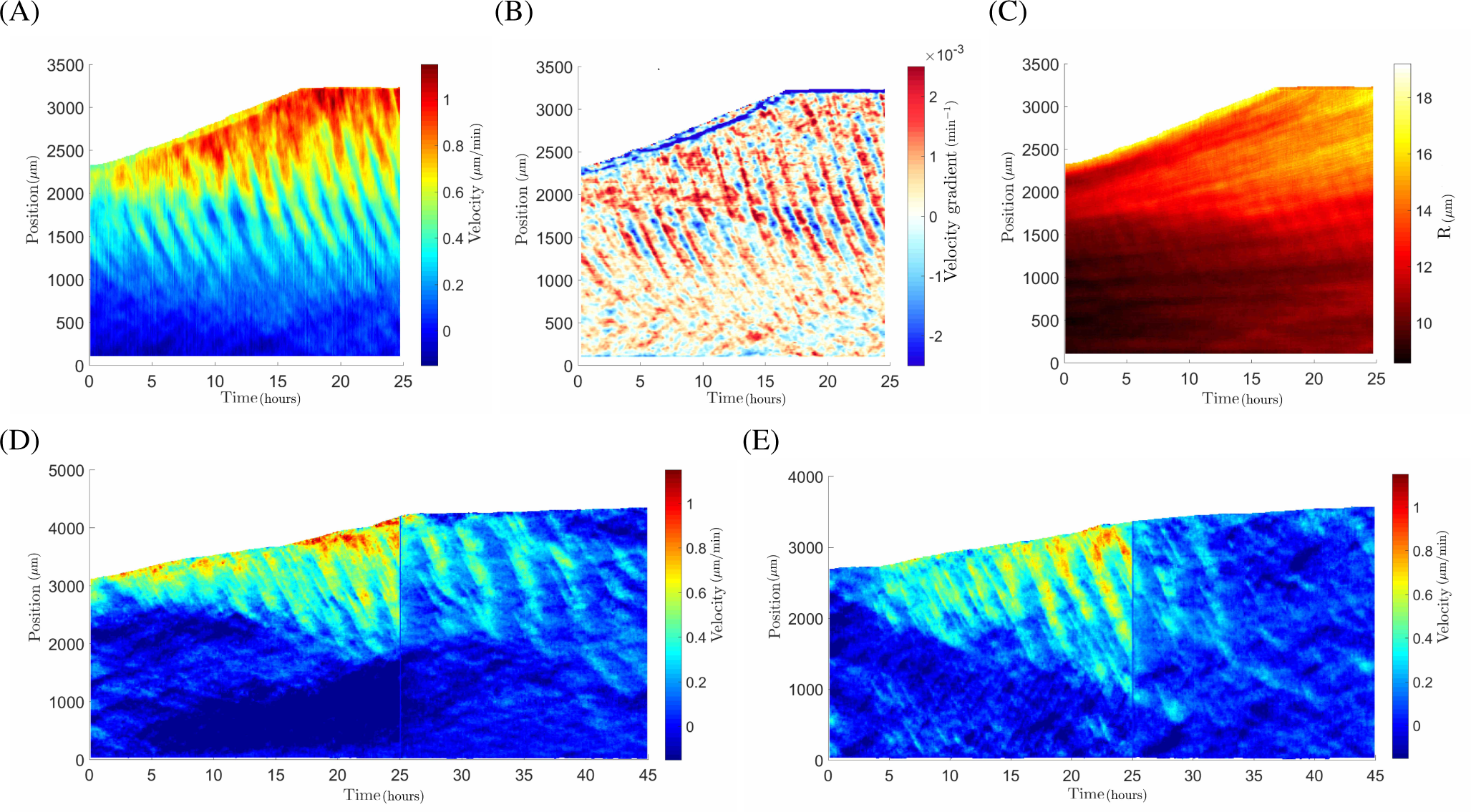
Propagating waves. (*A-C)* Space-time diagram (“kymograph”) of cell velocity, *V* (*A*), velocity gradient, 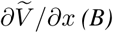, and effective cell radius, *R*(*C)*. Space *x* is from bottom (0 mm) to top (3 mm), time t from left (0 h) to right (25 h), and the top-left region is the bare substrate in front of the monolayer. All 8 strips showed similar results. (*D-E)* Same with application, after one day (visible as a vertical bar, since one image is not recorded), of CK-666 drug to inhibit lamellipodia formation, resulting in attenuation of waves (*D), N* = 7 strips; or in their almost complete suppression (*E), N* = 5 strips.

First measurements are manual. They indicate that the waves have a period around two hours, their wavelength is around one millimeter. Their amplitude decreases with the distance to the front: waves are not apparent at the other end of the monolayer, where the cell density is as high as 5 10^−3^ *μ*m^−2^ and the cell velocity vanishes. Where they are visible, their amplitude is steady in time, and large: it represents a relative variation in local velocity which typically ranges from 15% to 30% (Fig. S5B), i.e. up to 60 % crest-to-crest. Their velocity (indicated by the slope of the phase crests) is of order of ten micrometers per minute, and with a sign opposed to that of the cell velocity (indicated by the slope of the front position). Their phase crests are visibly curved: this evidences that the wave velocity (phase velocity) is larger near the front than in the middle of the monolayer.

Density small scale variations are dominated by local heterogeneities, which are signatures of initial density fluctuations (cells do not significantly mix nor rearrange) over a typical length scale of 200 *μ*m. More precisely, on the kymograph of *R* or 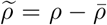 these fluctuations appear as bars which, near the front, are almost parallel to the front; far from the front, they are closer to horizontal (Fig. 2C, Fig. S6B); in between, along the line drawn on Fig. S6C which has a slope of 0.31 *μ*m/min, we measure 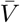 = 0.33 ± 0.02 *μ*m/min (SD). This proves that these fluctuations are advected at local velocity 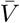 along with the monolayer itself. In addition, and although they are less visible, it is clearly possible to distinguish waves on the density (Figs. 2C, S5C, S6B), which have the same period as the velocity waves and are in phase opposition with them (Fig. S6C,D). They have a small amplitude, with a relative variation ranging from 1% to 2 % (Fig. S5D), i.e. up to 4 % crest-to-crest.

Inhibiting lamellipodia formation drastically decreases the monolayer average velocity (Fig. 1C, bottom; Fig. 2D,E). It also strongly decreases the amplitude of velocity waves (Fig. 2D), sometimes even almost completely suppressing them (Fig. 2E). The wave velocity, visible as the slope of the crests, is not significantly altered (Fig. 2D).

### Local wave characteristics

Phase crests are visibly curved (Figs. 2A, S6A): wave characteristics vary with space, and this can be quantified in several regions where the signal-to-noise ratio is sufficient (Fig. S5A). Using wavelets, we define, distinguish and measure locally 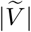 and *ϕ_V_* at each position *x* and time *t*, with 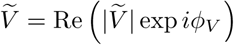, as follows.

The large space and time scales variations are encompassed by 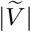; it tends to increase with 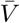 (Fig. S7A in the Supporting Material), and accordingly decrease with 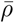. The small space and time scales variations are encompassed by *ϕ_V_* (Fig. S7B). The local phase *ϕ* of *V* in turn determines the local pulsation *ω* and wavenumber *k* according to the sign convention *ω* = *∂ϕ*/*∂t*, *k* = −*∂ϕ*/*∂x* (here *k* < 0); then the local time period *T* = 2π/*ω* and wavelength λ = 2π/|*k*|. While *k* varies in space (Figs. 2A, S6A), *ω* does not. Interestingly, comparing experiments performed at different densities shows that *ω* decreases with the front density (Fig. S3B). The wave velocity *c* = *ω*/*k* (here *c* < 0) is of order of minus ten times the cell velocity V; c does decrease with mean effective cell radius *R*_mean_ (Fig. S3C), i.e. |*c*| decreases with 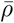. We do not observe that *ω*, *k* or *c* explicitly depend on the distance *X* to the front.

Again using wavelet analysis we define and measure 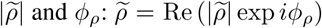. The kymograph of *ϕ_ρ_* shows that the wavelets detect the signal which physically corresponds to the wave velocity (i.e., wave velocity), and filter out irrelevent variations (Fig. S7B).

Inhibiting lamellipodia formation significantly decreases *ω* (Fig. S3B).

## Phenomenological description

Continuum mechanics (50) has been successfully used to model collective migration and wound closure of a cell monolayer on a substrate (51, 52). Several models have been proposed to explain the instability which gives rise to waves by invoking one of various active cell ingredients, within the constraints raised by symmetry considerations (11, 32, 53–57).

Building on these models, we propose a simple phenomenological description. We first recall cell number and momentum conservation laws within continuum mechanics, here expressed in one dimension. We then include an active force linked to cell polarization, to explain the existence of waves. Our goal is to perform testable predictions, compare them with experiments, and extract the values of relevant physical parameters.

### Conservation laws

In absence of cell division, the cell number conservation is expressed as

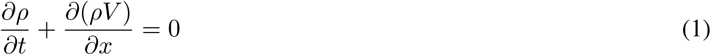

The local pulsation *ω*, wavenumber *k* < 0, and wave velocity *c* = *ω*/*k* < 0, measurable using wavelets, can be introduced here when we linearize Eq. (1) for small wave amplitude, i.e. neglecting the 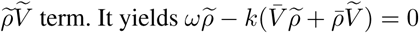, and:

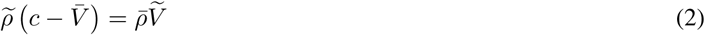

Since the order of magnitude of *c* is –10 *V*, Eq. (2) predicts that 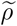 is in phase opposition with 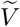,and that 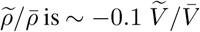. This explains why density oscillations are barely visible (Figs. 2C, S5C, S6B). By measuring the velocity and density wave characteristics at several points, we observe a local variability, which we exploit to check over a wide range that Eq. (2) is compatible with the observed oscillation amplitudes (Fig. S8A in the Supporting Material): Eq. (2) is checked with 10% precision. Phases even check Eq. (2) with 1% precision (Fig. S6D, Fig. S7B, Fig. S8B).

We now turn to the momentum conservation law, namely the force balance. The force equilibrium of the monolayer (integrated along the normal to the substrate) relates the external force per unit area *F*, exerted by the substrate on the cell monolayer, and the internal forces, namely the divergence of stress, as

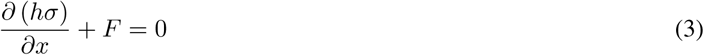

*σ* is the 1D stress (equivalent to the 3D stress component along *xx*) averaged over the monolayer thickness *h*. Note that this is a purely phenomenological description of stress, and in a real 2D description this would have to be replaced by the deviator of the stress tensor; alternative possibilities exist, such as the mean of the two principal stresses within the cell monolayer, i.e. half the trace of the stress tensor (32).

### Monolayer mechanical properties

In principle, the dissipation could be of both intra- or inter-cellular origin, and contribute to stress both in series or in parallel with elasticity (50). These different monolayer rheological properties are compatible with the appearance of waves (57), and it is beyond the scope of the present paper to enter such detailed description. To fix the ideas, the monolayer is often described as a viscoelastic liquid (11, 51), with a dissipative contribution in series with the elasticity (Maxwell model) and an elastic strain which relaxes over a viscoelastic time *τ*:

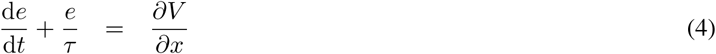

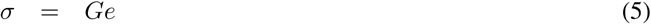

Here the the total strain rate is the velocity gradient *∂V/∂x* (Fig. 2B); the elastic strain rate is d*e*/d*t* = ∂*e*/∂*t* + *V∂e*/*∂x*; *G* is the elastic modulus typically in the range 10^2^ – 10^3^ Pa, obtained for single cell (58), by micro-indentation (59, 60) or on a monolayer (61) (note that stretching a suspended monolayer, including cell-cell junctions, yields a much larger value ∼ 2 10^4^ Pa (12)); the value of *τ* is discussed below. From these orders of magnitude, we predict that waves of traction force, if they exist, are below our current detection limit.

### Active mechanical ingredients

In the literature, the force per unit area exerted by the substrate on the monolayer is often expressed as the sum of active and friction contributions, *F* = *f_a_p* – *ζV* (14). Here *f_a_* is the characteristic value of the active cell force a cell can exert; it is of order of 300 Pa (5)), and decreases with *ρ* (7, 8, 13). The dimensionless real number *p* is the one-dimensional equivalent of the cell polarization. It is convenient to introduce *V_a_* = *f_a_*/*ζ* which corresponds to a characteristic active migration velocity, and write

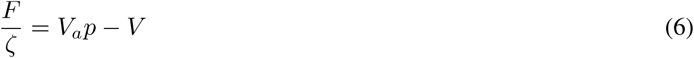

Note that, alternatively, it would have been possible to consider *p* as a Boolean variable, being either +1 or −1, while *V_a_* and *f_a_* are continuous variables. This alternative could be important when discussing for instance how the polarity *p* is related with biochemistry, and whether it could change sign by passing continuously through 0; but such debate is beyond the scope of the present paper.

We use the values of the friction coefficient *ζ* ∼ 10^9^ N m^−3^ s (14, 32). For a wavenumber *k* ∼ 10^4^ rad m^−1^, and with upper estimates of the 3D cell viscosity *η* of order of 10^2^ Pa s (62, 63), we obtain that the modulus of the internal viscous force *ηk*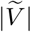 is at least thousand times smaller than that of the typical external friction force, *ζ*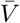. We thus neglect the viscosity contribution in parallel with the elasticity (32).

Developing Eq. (1) lets a spatial derivative of *V* appear. Injecting it in Eq. (6) involves a derivative of *F*, then Eq. (3) a second derivative of *σ*, and finally Eq. (5) a second derivative of *e*:

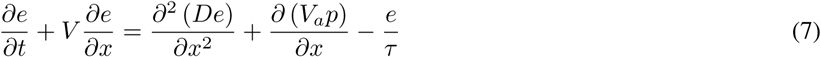

Since *h* varies slowly with space, Eq. (7) is locally a diffusion equation (64), with an effective diffusion coefficient *D* = *Gh*/*ζ*; *D* increases with monolayer stiffness and decreases with friction. In principle, the steady flow can become instable, and waves appear, if the heterogeneity of the active term *V_a_p* overcomes the stabilising diffusion term. A heterogeneity in migration force might create a heterogeneity in velocity, affecting in turn the stress, which would feed back on the force. The question is how this feedback could become positive.

### Coupled polarity equation

Polarity can couple to cell strain through a mechano-sensitive protein or protein complex (such as Merlin (65)). Here we assume that the monolayer is already polarized, symmetry is broken due to the propagating front (hence symmetry constraints (56) are not enforced in the following equations). Cells already have a polarity *p*, which is enhanced by cells stretching and decreased by cell compression. We neglect: non-linearities; viscosity; and the large scale variation of *ρ* and 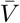 over the whole strip length scale. These simplifying hypotheses can easily be relaxed if required, for instance if future experiments add new details. We have checked *a posteriori* that a complete treatment which includes the space variations of *ρ* and 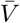 increases the present predictions of |*c*| and *ω* by less than 10 %.

We study the stability of a homogeneous steady state where all cell migrate in the same direction, 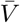 = *V_a_* > 0, and are positively polarized, *p̅* = 1. The density is 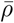, the traction force 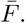 We study for instance the case 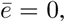 which is relevant in the region close to the front where the waves are most visible, and which is the value towards which *e* relaxes when *τ* is finite. Note that the symmetric steady state, with 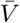 and 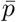 negative, is irrelevant here (unlike in symmetric migration experiments (11)). The third steady state, 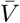 = 0, exists initially, but ceases to be stable when the confinement is removed, and is also irrelevant here.

The polarity follows the strain with a delay, *τ_ρ_*, which in two or more dimensions would be the reorientation time (51):

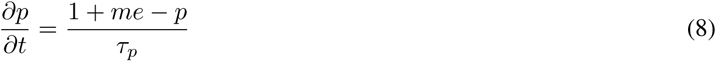

Here *m* is a non-dimensional factor coupling between polarity and cell strain, and when *me* is of order 1 the polarity value changes by one unit. If we integrate Eq. (4) in time, using the observed velocity wave, we observe that the total strain has an amplitude of order 0.1. It means that the elastic strain has an amplitude of at most 0.1, and probably around half of it if de/dt and *e*/*τ* are comparable (which is the case since *τ* is comparable with the wave period, see below). Hence a strain of at most 0.1 (and possibly 0.05) suffices to change the polarity from value 0 to value 1, and m is of order of 10, possibly 20, or 25 at most.

### Onset of wave appearance

To perform a linear stability analysis around the steady state *e* = 0, *p* = 1, the small variables are *δe* = *e* and *δp* = *p* − 1. Calculations near *ē* = 0 are simple, as terms of order *ē* are suppressed, and as terms due to variations of h are second order and thus negligible. Eqs. (7),(8) become, after linearization:

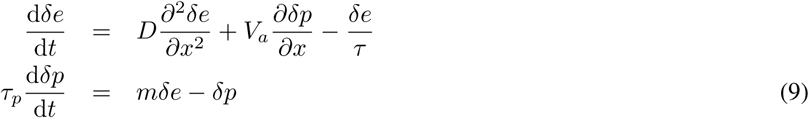

We look for small perturbations (of the steady, homogeneous state) proportional to exp (*st* – *ikx*), where the wavenumber *k* is a real number, and the wave growth rate s is a complex number with a real part which is strictly positive when the steady state is unstable, Re(*s*) > 0, and a non zero imaginary part, Im(*s*) ≠ 0. The Jacobian matrix of the equation system, Eqs. (9), is:

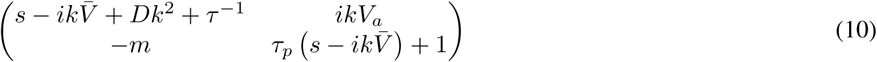

Solving in *s* simply requires to write that the determinant of 𝒥 is zero:

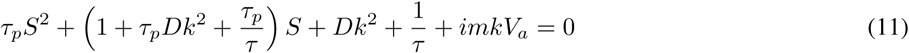

where 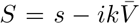. There are two roots, which depend on *k* and on parameter values. We numerically solve Eq. (11) and look for a root with Re(*s*) > 0 and Im(*s*) ≠ 0. Depending on the parameter values (Fig. 3), there exists a range of *k* with one such root, and the steady, homogeneous solution is unstable. A propagating wave appears; the mode k which develops more quickly is that for which Re(*s*) is maximum, i.e. dRe(*s*)/d*k* = 0 (until the amplitude increases enough to reach the non-linear regime). Its imaginary part Im(*s*) is the pulsation *ω* of this mode.

**Figure 3:**
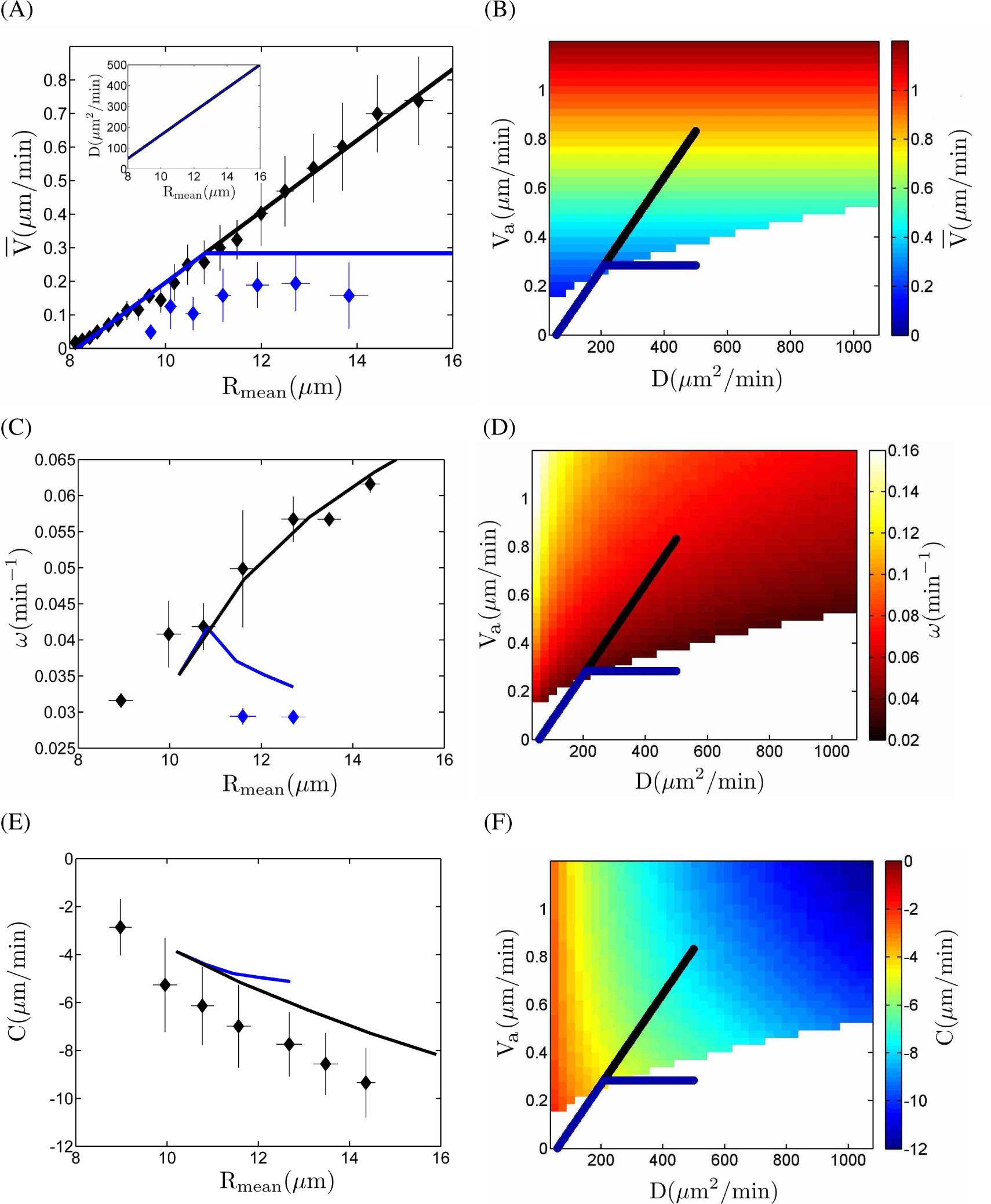
Predictions of phenomenological model. Parameter values (*m* = 25, *τ_p_* = 15 min, *τ* = 180 min) are manually chosen to obtain a good agreement with the data, see text. *(A)* 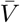 vs *R*_mean_. Points are experimental data (Fig. 1C, bottom). Black: linear relation, for experiments with mitomycin. Blue: estimated relation with saturation, for experiments with lamellipodia inhibition using CK666. Inset: *D* vs *R*_mean_, estimated relation, not affected by CK666. *(B)* Diagram of cell velocity 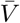 vs (*D*, *V_a_*). It is plotted in the region where the wave amplitude growth rate is positive (existence of waves). The regions where the wave amplitude growth rate is negative (the steady migration is stable) are left blank. Lines correspond to those in (A); *R*_mean_ is increasing from bottom left to top right. *(C, D)* Same for wave pulsation *ω*. *(E, F)* Same for wave velocity c; note that its values are negative, and that with CK666 the values of *c* are too noisy for quantitative measurements, but are similar.

To fix the ideas and provide example of calculations, we take from experiments that 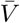 is of order of a *μ*m.min^−1^. The delay time *τ_p_* of polarity with respect to stretching, due to the reaction time of Rac, could be of order of 25 min (65). The value of *m* also affects the predictions; higher *m* values tend to make the absolute value of *c* larger (*c* is more negative). We now discuss the value of *τ* and defer the discussion of *V_a_* and *D* to the next section.

In the viscoelastic liquid description we consider here, at time scales smaller than *τ* the monolayer can sustain a shear stress and behaves as mostly solid, while at longer time scales the strain relaxes and the monolayer behaves as mostly liquid. So it is important to determine whether *τ* is larger or smaller than the time scale of the waves, of order of one hour. This is subject to debate, since depending on the cell line the elastic modulus and the viscoelastic time of tissues can vary over orders of magnitude (66). Even when restricting to MDCK monolayers, published values for the viscoelastic time *τ* range from 15 min (51) to 3 - 5 hours (62, 66). Several articles (5, 11, 52) choose to treat the monolayer as elastic, given that the elastic modulus can be an effective modulus arising from the cell activity (66, 67).

We have checked numerically that small values of *τ*, for instance 30 min or less, stabilize the steady state, while large values of *τ*, 1 hour or more, allow for the wave appearance. The wave characteristics we determine barely change when *τ* spans the range 1 to 5 hours.

### Wave characteristics

*V_a_* makes waves appear while *D* is damping them. Experimental measurements of wave characteristics can help estimate orders of magnitude of *D* and *V_a_*. However, determining their precise values is difficult and strongly dependent on the model (which is here only phenomenological). We thus let values of *D* and *V_a_* vary within a reasonable range and solve systematically Eq. (11), to determine a phase diagram in the (*D*, *V_a_*) plane.

In Fig. 3 we plot the model predictions for parameter values *τ_p_* = 15 min, *τ* = 180 min, which are realistic (see above), and *m* = 25, chosen to obtain a good agreement with the experimental data. These data indicate that both *V_a_* and |*c*| increase linearly with R_mean_ (Fig. 1C, bottom; Fig. 3E, Fig. S3C). We use the measured relation of 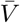 = 0.106 *R*_mean_ – 0.864 (Fig. 1C, bottom, inset; Fig. 3A). To reproduce the experimentally observed linear variation of |*c*| with *R*_mean_ (Fig. 3E), we find that *D* too has to increase with *R*_mean_; in the following we choose a linear relation between *D* and *R*_mean_ (inset of Fig. 3A). This determines the black lines on Fig. 3B-F as possible variations of 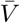, *c* and *ω*, and corresponding trajectories in the (*D, V_a_*) plane, when *R*_mean_ varies. We obtain an agreement with measurements of *ω* (*R*_mean_), which we reproduce quantitatively, and of *c*(*R*_mean_), which we capture qualitatively. Note that *c* is negative, indicating backwards waves as observed in experiments.

The instability threshold, visible as the limit between colored and blank regions (Fig. 3B,D,F), indicates when *V_a_* is strong enough to overcome *D*. We obtain a consistent pictures with typically *D* of order of 10^2^ *μ*m^2^/min (i.e. *G* of a few 10^2^ Pa), *V_a_* of order of 10^−1^ *μ*m/min (*f_a_* of one or a few 10 Pa), wavelength of several 10^2^ or 10^3^ *μ*m, time period of a few 10^2^ min, |*c*| of order of 10^1^ *μ*m/min.

We observe that CK666 results in a saturation in 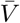, probably linked with a limit in lamellipodia size (43) (Fig. 1C, bottom). Since within the model *V_a_* = 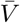 we use at small *R*_mean_ values the same linear increase as without drug (Figs. 1C, bottom, inset; Fig. 3A), then a plateau. To determine the position of the cross-over between these regimes, i.e. the onset of saturation, we observe that with CK666 the monolayer is very close to the limit of wave appearance: waves are either decreased in amplitude, and slowed down, or suppressed. We thus position the cross-over at the intersection between the straight line and the instability threshold.

Experimentally, in the case where small waves are observed in presence of CK666, then *R*_mean_ is near 12 *μ*m, *ω* is near 0.03 min^−1^. In this case the wave velocity *c* measurement is too noisy to be quantitative, but *c* seems unaffected as no slope rupture is visible on the phase crests (Fig. 2D). These features are qualitatively captured by our model.

## Discussion

### Cell migration results

Experiments had shown that *V* decreased with the distance *X* to the moving front (11) while *ρ* and *σ_xx_* increased with *X* and were proportional to each other (5). We can imagine two possible interpretations: either that both *ρ* and *σ_xx_* happen to vary similarly with *X*, with a reinforcement of cell-cell junctions from the front to the back; or that *σ_xx_* is actually determined by *ρ*.

Here, we observe a large enough range of cell densities and velocity, with a good enough signal-to-noise ratio, to discriminate between dependence with *X* vs with *ρ*. We find that in the monolayer bulk, *V* depends only on *ρ*, namely that it increases linearly with *R*_mean_, irrespectively of *X* or of the past history of the cell monolayer (advection, divisions, extrusions) which causes the observed density value. This is compatible with observations of Refs. (5, 11); it suggests that the traction force is cell-autonomous and is linear in *R*_mean_; and it favors our above interpretation that *σ_xx_* is determined by *ρ*.

With migration alone, coherent cell collective movement propagates over a few mm inside the monolayer. The wave wavelength is of order of 1 mm and more broadly speaking the whole velocity field is established coherently over 4 mm. We have used 200 *μ*m and 1 mm wide strips, larger than the typical 200 *μ*m correlation length for the cell velocity field (9, 28). We do not detect any significant effect on the strip width on the results presented here.

Our observations are compatible with a large-scale polarized activity induced for instance by activity of the protein Merlin (a tumor suppressor), as recently shown experimentally (65). The cells at the front of the migrating monolayers are known to exert large traction forces (61, 68) that can induce the build-up of a large intercellular stress and in turn, a polarization of the following cell by a relocalization of Merlin from the cell-cell junctions to the cytoplasm. When the cell is at rest, Merlin is localized at the cell-cell junctions. This junctional Merlin inhibits the formation of cryptic lamellipodia. On the other hand, when cell-cell junctions experience a stretching stress, Merlin is relocated to the cytoplasm. Due to the decrease in junctional Merlin, the Rac pathway becomes activated, within a delay time of a few tens of minutes. Then, within a much smaller delay, Rac activates the generation of cell polarization and lamellipodia, responsible for the migrating forces (65). The iteration of such processes may lead to large scale polarization within the tissue.

The most visible effect of divisions (Fig. 1C, bottom; Fig. S4) is to steadily increase the density, which contributes to jamming and decreases the migration velocity. However, at given density, the divisions themselves contribute to increase the monolayer movement and the front velocity. In addition, divisions increase the noise in cell velocity; more regions have a negative velocity (Fig. S4D).

To complement existing experiments with blebbistatin which focus on the role of cell contractility (11, 32), we inhibit lamellipodia with CK666. It has a clear effect, even in the bulk of the monolayer, on *V* which is decreased; and on the *V*(*R*_mean_) relation, which saturates and is no longer linear (Fig. 1C, bottom). This suggests that the contribution of lamellipodia to the traction force is dominant, and linear in *R*_mean_.

### Wave results

In experiments with divisions, the mechanical waves are slightly visible and overdamped; this is broadly compatible with the literature (11, 30–32). Without divisions, we obtain a good enough signal-to-noise ratio to measure the wave properties. Moreover, thanks to this signal-to-noise ratio and experiment reproducibility, we can even measure the variation of wave properties across space and time, and with enough precision to discriminate between dependence with position vs with density. We observe that the wave velocity *c* is ∼ –10 *V* and, like *V*, it depends explicitly only on *ρ*: it is linear in *R*_mean_, again irrespectively of *X* or of the past history of the cell monolayer.

For a given experiment, although *ρ* is space-dependent, the wave pulsation *ω* is spatially homogeneous. It might arise from the fact that the most developed mode temporally forces the instability over the whole monolayer, resulting in a space-dependent wavenumber *k*. Now, comparing experiments at different densities, we observe that *ω* increases with *R*_mean_ (Fig. 3C, Fig. S3B). Inhibiting the lamellipodia decreases *ω*, at a given *R*_mean_.

Backwards-propagating waves are reminiscent of a generic instability mechanism originally discussed in the context of car traffic (69), which arises when the velocity *V* is a decreasing function of density *ρ*. For instance, velocity pulses have been observed for dense colloids in channel flow near jamming - unjamming transitions, in experiments (70) and in simulations (71). Similarly, self-propelled agents, which tend to accumulate where they move more slowly and/or slow down at high density (for either biochemical or steric reasons) undergo a positive feedback which can lead to motility-induced phase separation between dense and dilute fluid phases (72, 73).

### Comparison of experiments with phenomenology

Inspired by published observations and by our own, we propose here a simple description where motility forces in the bulk of a homogeneous monolayer are oriented by a dynamic pulling on cell-cell junctions. This elasticity-polarity coupling is injected in classical rheology equations of continuum mechanics.

Waves spontaneously appear close to the instability limit (Fig. 3). This could explain why waves had been difficult to detect in the past (11, 30) (see also the Supp. Figs. S2, S5 of Ref. (32)), as parameters can change depending on cell size or substrate coating. With reasonable parameters values, typically *G* of a few 10^2^ Pa, *f_a_* of one or a few 10 Pa, we find propagating waves with wavelength of several 10^2^ or 10^3^ *μ*m, time period of a few 10^2^ min, of the same order as the experimental values. We predict a negative wave velocity *c* indicating backwards waves, with |*c*| of order of 10^1^ *μ*m/min, as observed in experiments. This is probably because a cell migrating towards the front pulls the cell behind it and favors its migration (in our model, under traction the cell polarity increases, with a positive coupling factor *m*).

We expect that when *ρ* decreases and *R*_mean_ increases, *V_a_* increases significantly (Fig. 3). It is compatible with our observation that, when comparing experiments at different densities, *ω* decreases with *ρ* (Fig. 3C). Our model presents a Hopf bifurcation sensitive to the density, and a slowing down of the wave pulsation when approaching the bifurcation. With proliferation due to cell division, the density increases; it can lead either to decrease in wave amplitude and pulsation, or to a suppression of the wave, as observed in experiments (Fig. 2D,E, Fig. 3C). The model suggests that two experiments performed at a slightly different initial density can, after lamellipodia inhibition, lead either to decrease in wave amplitude and pulsation, or to a suppression of the waves; this could explain the observed effects of CK666 (Fig. 2D,E, Fig. 3C).

Our model qualitatively suggests that *D* increases with *R*_mean_ (Fig. 3A, inset). Our estimation of *D* which increases linearly, by a factor of ten, when *R*_mean_ doubles (Fig. 3A, inset) is compatible with the observation that the elastic modulus can vary over orders of magnitude, and scales linearly with the size of the constituent cells (66). This could be compatible with the intuition that a cell which spreads has more stress fibers and a more organised cortex, resulting in a larger cell stiffness *G*. It is also compatible with the fact that the relation *D* (*R*_mean_) is much less influenced than the relation *V_a_* (*R*_mean_) by the lamellipodia inhibition. Note that alternative explanations exist, for instance if the friction coefficient *ζ* was decreasing with *R*_mean_.

In summary, our model (Fig. 3) is precisely compatible with measurements of 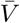, reproduces quantitatively *ω* and qualitatively *c*. Our predictions of how *V_a_* and *D* vary with *R*_mean_ agree with independent experiments on the same MDCK cells (66). In presence of CK666, our model is qualitatively compatible with either the suppression of waves, or waves with a smaller *ω* and an unchanged *c*.

### Comparison with existing models

Trepat and coworkers present numerical simulations (11) where, based on their cell stretching experiments, they introduce a dynamically changing elasticity (non-linear cytoskeletal reinforcement). Our ingredients are similar to theirs except that, inspired by recent experiments on collective cell migration (65), we introduce a dynamically changing traction force modulated by the elastic strain. Note that their simulations predict forward-propagating waves, i.e. waves velocity in the same direction as cell velocity. On the other hand, their observations (11, 29), our observations and our model agree that waves propagate backwards. We are not aware of any interpretation of this discrepancy.

Blanch-Mercader and Casademunt (55) too predict forward propagating waves. They explain that even viscous tissues can have an effective elasticity in which waves appear. This is compatible with the idea, arising from both modeling (67) and experiments (66), that the elastic modulus can be an effective modulus arising from the cell activity.

Marcq and co-workers have shown that since cells are active, the appearance of waves is compatible with a wide range of ingredients. In particular, the rheology is not crucial, as waves can appear in materials with various rheological behaviours (54, 57); this is compatible with the conclusions of Blanch-Mercader and Casademunt (55). They predict that there are more waves when divisions are inhibited (56) and less waves when lamellipodia are inhibited (57). Our observations and model agree with these predictions.

Banerjee, Marchetti and coworkers (32, 53) predict the existence of waves depending on an effective elasticity and traction force amplitude. Our model is based on an approach similar to theirs; we make it simpler while keeping the main ingredients. They find a wave velocity comparable to a cell length divided by the time (∼ 2 min) required for mechanical stress information to propagate across the cell. Their wave period, around 6±2 h, increases with decreasing traction forces. Their predictions are consistent with three of our observations. First, their waves propagate backwards. Second, the wave temporal period decreases with increasing density (and thus decreasing traction force) at the migrating front. Third, when adding CK666, either the wave temporal period increases or the wave disappears.

In summary, our experimental observations and quantitative characterization of backwards propagating velocity wave characteristics, and their dependence in the model parameters, should strongly constrain analytical models or numerical simulations in the future. Our predictions can be tested experimentally. Such studies should shed light on the role of long-range information propagation mediated by mechanical stress, during both *in vitro* cell migration and *in vivo* embryogenesis.

## Author contributions

S.T., B.La., H.D.-A., F.G. designed the research; S.T., E.G., performed experiments; S.T., E.G., B.Li, O.C., analysed experiments; S.T., B.Li, F.G. developed the model; S.T., B.Li, H.D.-A., F.G. wrote the manuscript; all authors discussed the results and the manuscript.

## Acknowledgments

We warmly thank P. Marcq, S. Yabunaka for critical reading of the manuscript, and T. Das, F. Gallet, Y. Jiang, U. Schwarz for discussions. This work was supported by the European Research Council under the European Union’s Seventh Framework Program /ERC consolidator grant agreement 617233 and the LABEX “Who am I?”.

## Supporting Information

**Supporting Figure S1.**
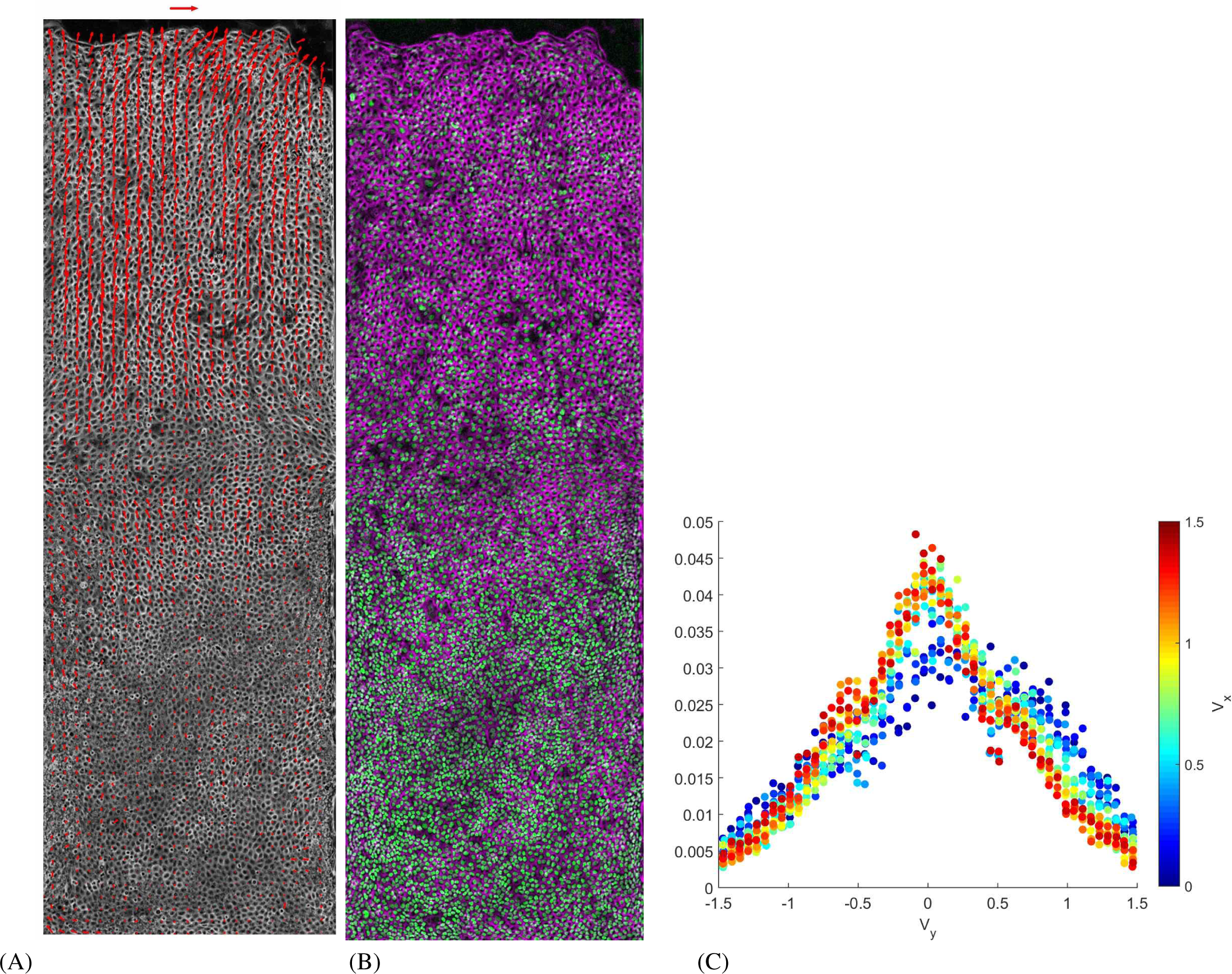
Cell velocity and density. *(A)* Instantaneous velocity field 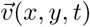 superimposed on cell contours, at *t* = 15 h after the first image (i.e. after around 20 h of migration). Scale arrow: 2 *μ*m/min. *(B)* Corresponding picture of nuclei labeled with histone GFP (green) evidence the large scale cell density variation; they can be detected for cell density measurement, and tracked for cell velocity validation. Cell contours appear in purple. *(C)* Distributions of *y* component of velocity field 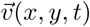, while its *X* component is indicated by the color code.

**Supporting Figure S2.**
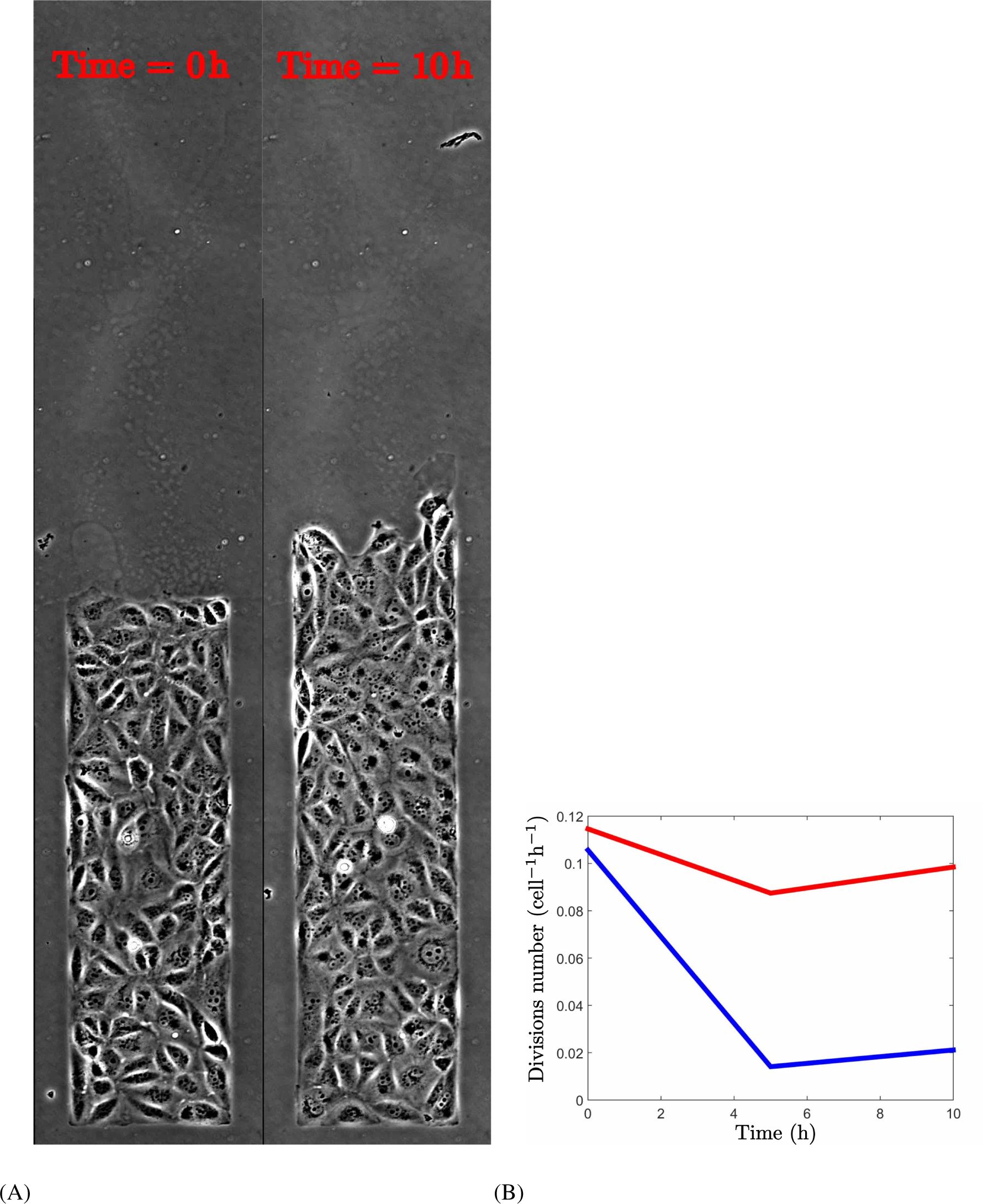
Effect of mitomycin. *(A)* First (left) and last (right) images used to test the effect of mitomycin during 10 hours after rinsing (*t* = 0). Strip width 200 *μ*m. A small length is chosen, so that the whole experiment is visible on each image. *(B)* Average division rate per cell and per hour. Blue: with mitomycin, rinsed at *t* = 0. Red: control, no mitomycin.

**Supporting Figure S3.**
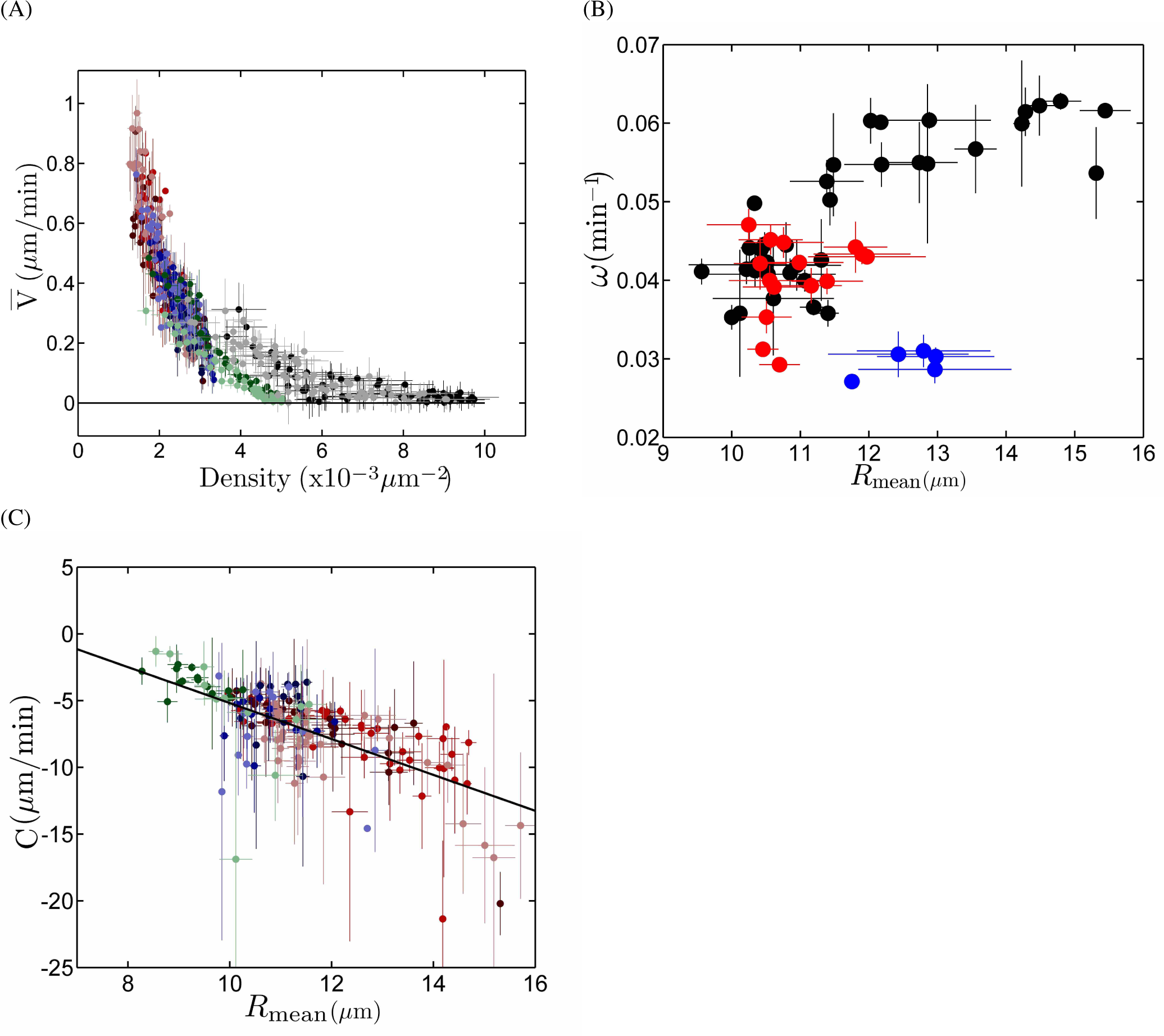
Variations with cell density. (A) Cell velocity 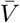 vs cell density 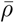. Same color code as in Fig. 1C. *N* = 319 with mitomycin C, colors; *N* = 134 in standard conditions, grey levels. (B) Pulsation *ω* determined by wavelet analysis in the region of interest (which is manually selected according to its good signal-to-noise ratio, Fig. S5A), vs front value of *R*_mean_. Each point results from the spatial average ± SD both of *ω* and of *R*_mean_, over a large spatial box near the front, at a same time (binning 528 *μ*m *X* 180 min). Black: with mitomycin C; experiments as in Fig. 2A, *N* = 8 strips. Red: experiments as in (Fig. 2D, before adding CK666; *N* = 5 strips. Blue, same after adding CK666 (points are measured at later times, and thus lower *R*_mean_); *N* = 3 strips. Strips with signal-to-noise ratio too low to be measurable are not plotted. (C) Wave velocity *c*, determined by wavelet analysis in the same region of interest, versus mean effective cell radius *R*_mean_, with a linear fit *c* = 9.2 – 1.4 Rmean (R=0.59); *N* = 155 data points (one outlier has been removed).

**Supporting Figure S4.**
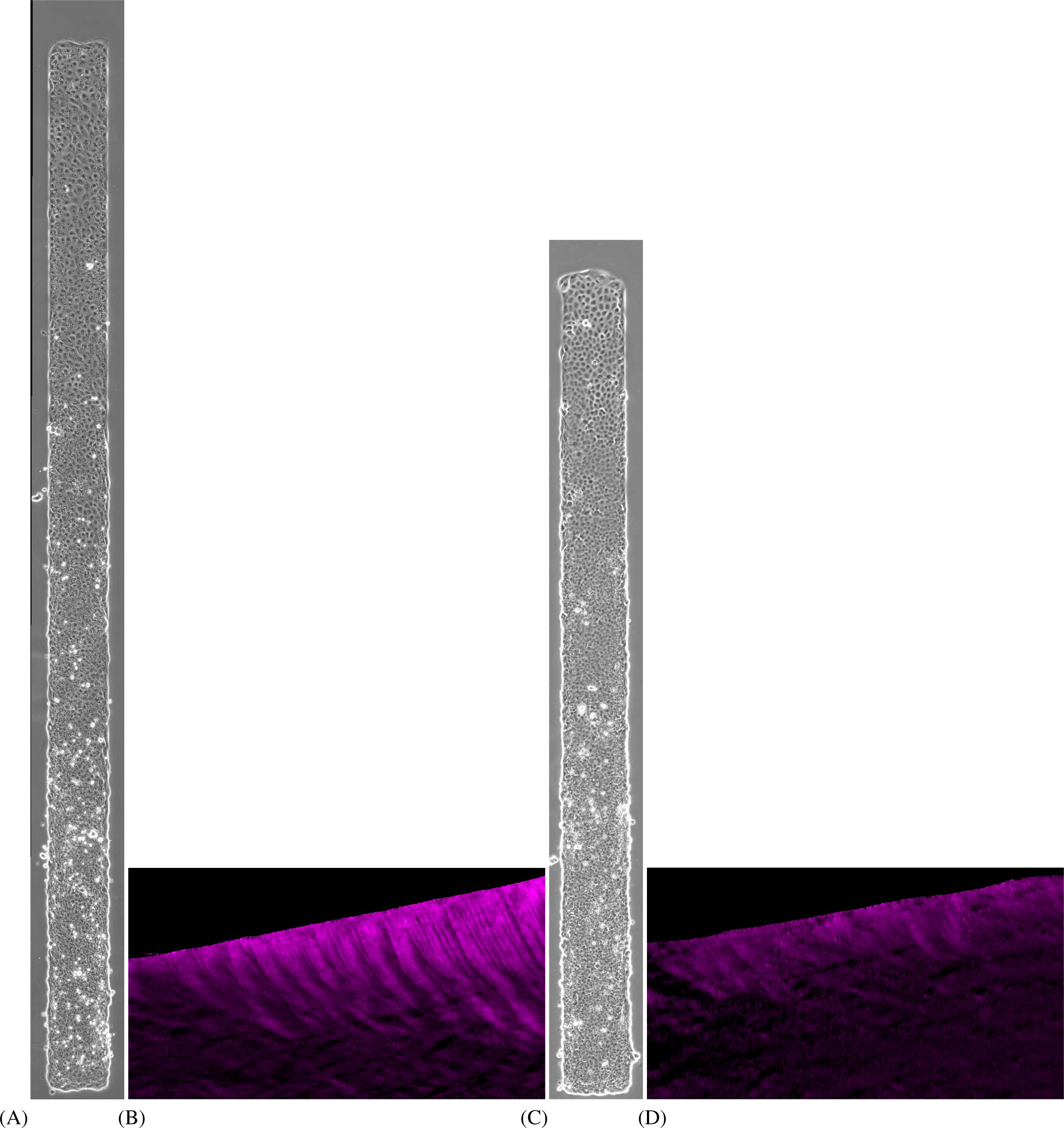
Effect of divisions, in small width strips. (A) Experiment performed with mitomycin C to prevent divisions. Phase-contrast images showing cell contours. (B) Corresponding kymograph of cell velocity *V*(*x,t*) averaged over *y* but not over time. *(C,D)* Same under standard conditions (i.e. without mitomycin C). In *(A,C):* Strip width 200 *μ*m; migration from left to right; substrate: rigid PDMS. Despite the difference in the initial cell density, 10 10^−3^ *μ*m^−2^ *(A)* vs 5 10^−3^ *μ*m^−2^ *(C)*, at the time the picture is taken (*t* = 30 h) the density ranges in both experiments overlap. In *(B,D)*: Space *X* is from bottom (0 mm) to top (3.8 mm in B, 2.7 mm in D), time *t* from left (0 h) to right (30 h), and the top-left region is the bare substrate in front of the monolayer. Color code *V* from −0.15 (black) to 1.15 *μ*m/min (purple).

**Supporting Figure S5.**
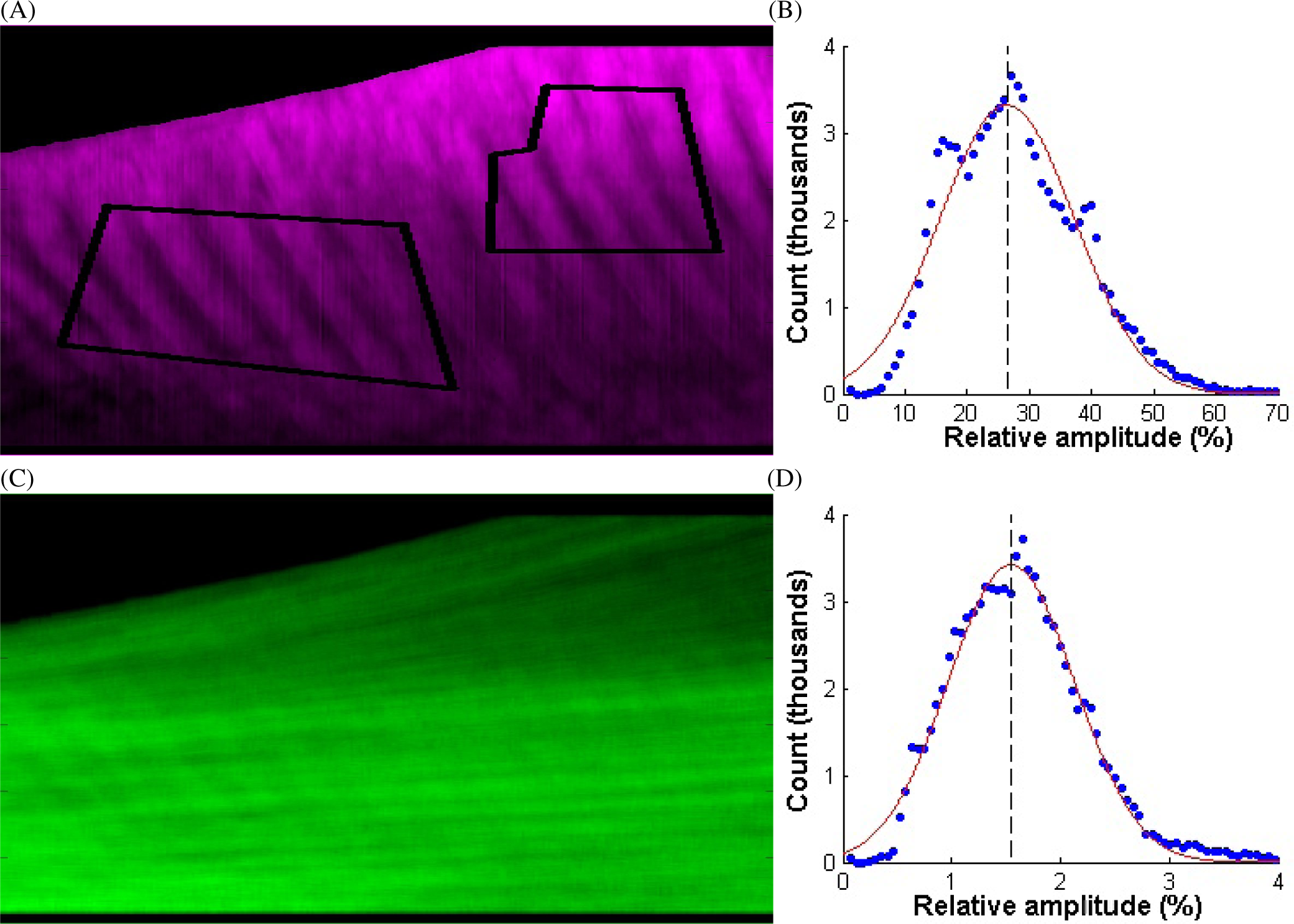
Local spatial variations. *(A)* Cell velocity *V*(*x, t*) averaged over *y* but not over time. Space *X* is from bottom (0 mm) to top (3 mm), time t from left (0 h) to right (25 h), and the top-left region is the bare substrate in front of the monolayer. Color code 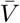 from —0.15 (black) to 1.15 *μ*m/min (purple), 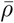 from 0.5 10^−3^ *μ*m^−2^ (black) to 3.5 10^−3^ *μ*m^−2^ (green). The outlined region of interest is selected manually as the region where waves have a good signal-to-noise ratio, large enough to perform the wavelet analysis. The results presented here are robust with respect to this manual selection. *(B)* Histogram of relative amplitude of wave velocity cumulated for 8 different experiments. Red line: Gaussian fit, of average 26.4% (marked by black dashes) and width 15.3% *(C,D)* Same for density. Gaussian fit: average 1.55% and width 0.83%.

**Supporting Figure S6.**
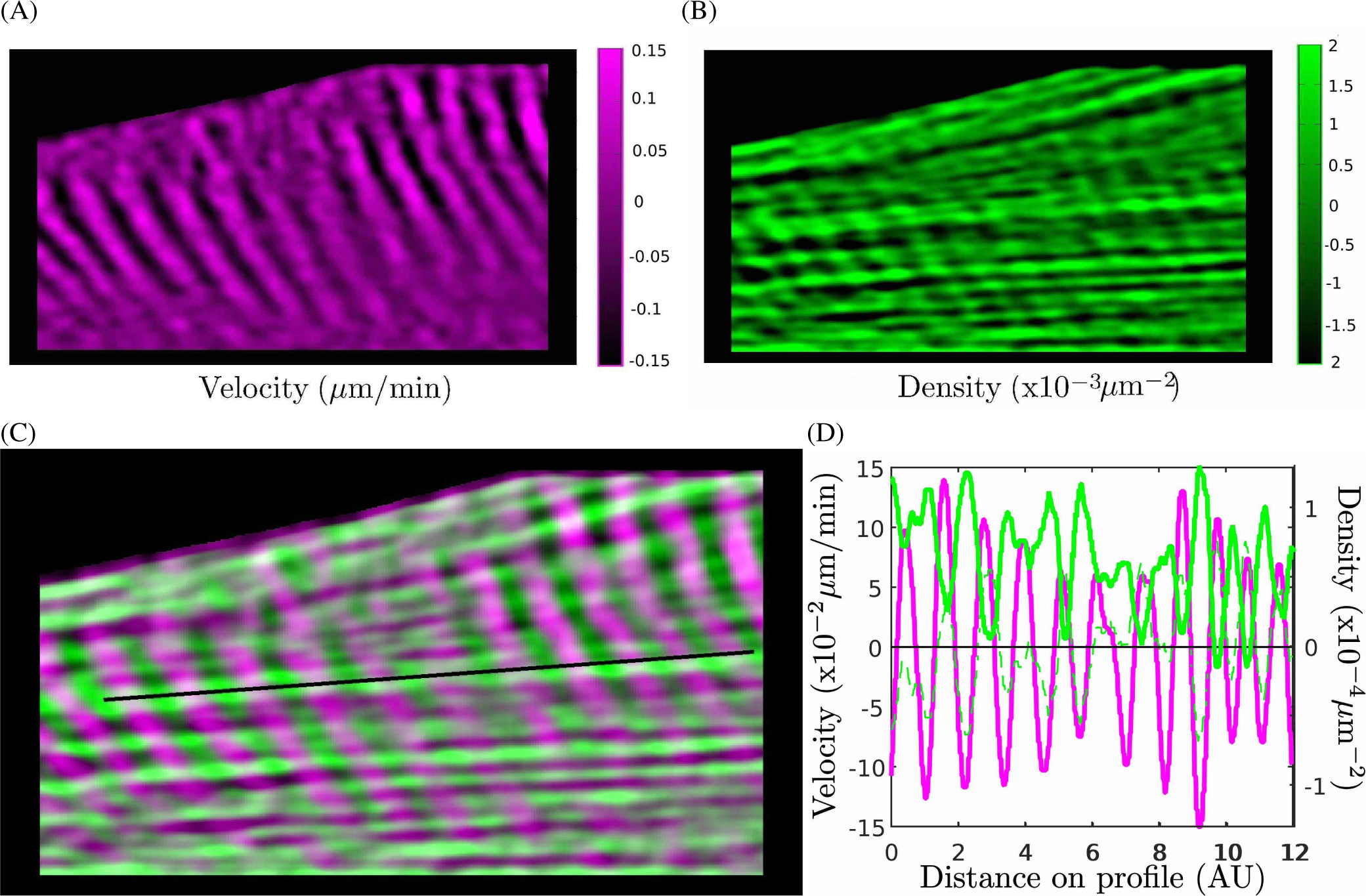
Propagating waves. *(A-C)* Kymograph of small scale variations in cell velocity, 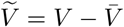 *(A)*, in density, 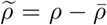 *(B)*, and merge, *(C)*. Space *x* is from bottom (0 mm) to top (3 mm), time *t* from left (0 h) to right (25 h), and the top-left region is the bare substrate in front of the monolayer. Color code 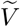 from −0.15 *μ*m/min (black) to 0.15 *μ*m/min (purple), 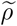 from —1.3 10^−3^ *μ*m^−2^ (black) to 1.3 10^−3^ *μ*m^−2^ (green). The line is manually drawn along a density local maximum. *(D)* Plot of 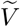 (purple) and 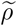 (green) along the line manually drawn in *(C)*. Inverting the sign of 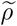 (green dashes) evidences its phase opposition with 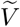, which agrees with Eq. (2).

**Supporting Figure S7.**
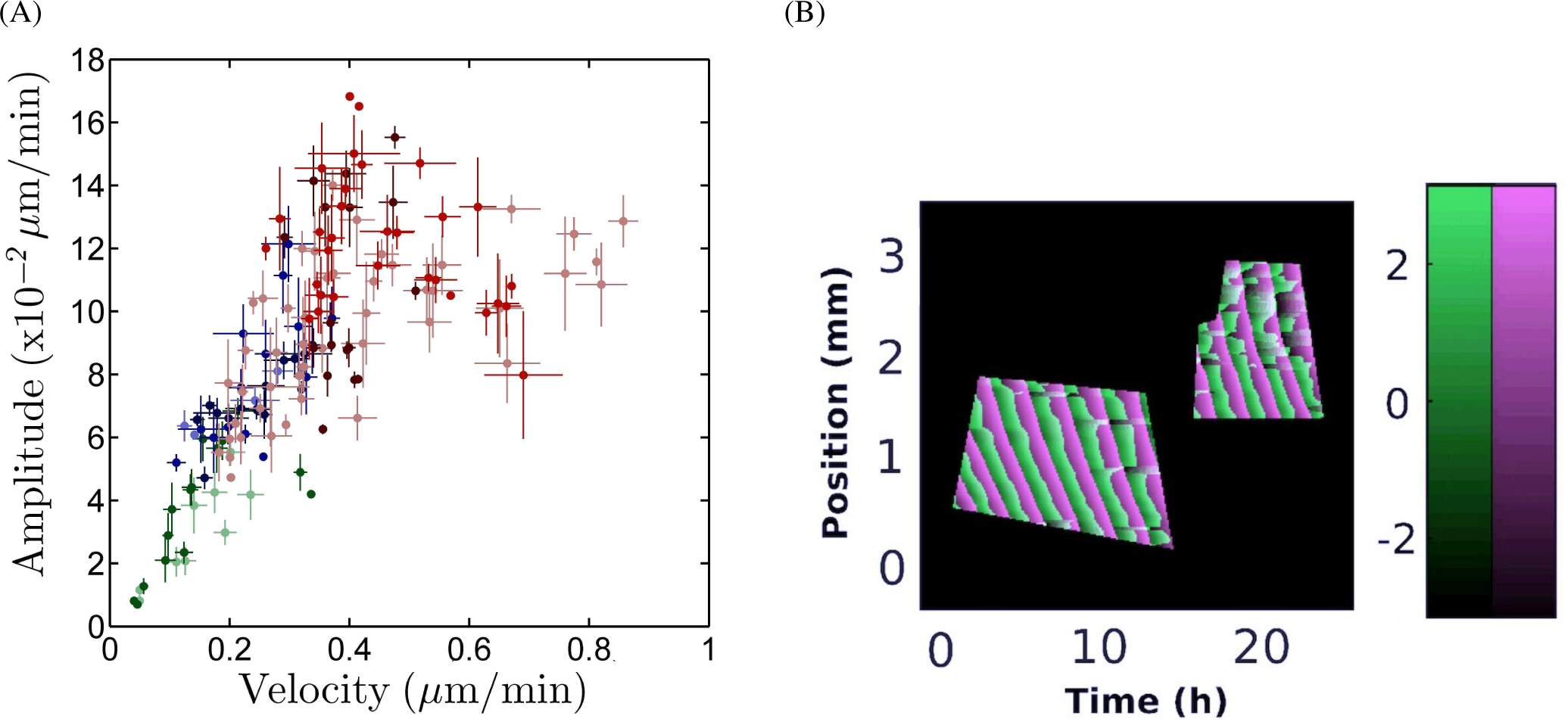
Local wave characteristics. They are determined by wavelet analysis in the region of interest, which is manually selected according to its good signal-to-noise ratio (Fig. S5A). *(A)* Amplitude 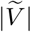 (± SD) of velocity waves, versus average cell velocity 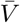; *N* = 156 data points. Same color code as in Fig. 1C. *(B)* Merged kymographs of phases: *ϕ_ρ_* from −π (black) to π (green), and *ϕν* from –π (black) to π (purple).

**Supporting Figure S8.**
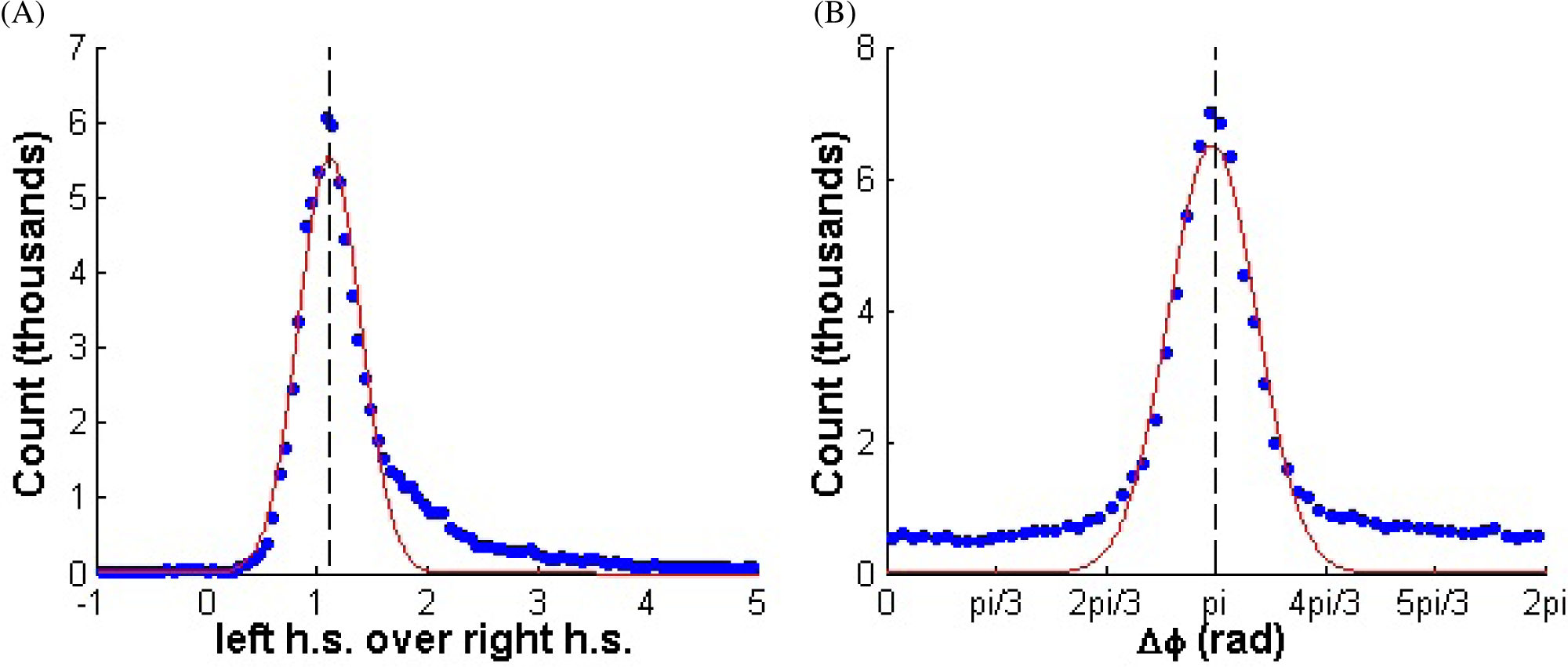
Test of equation for density. (A) Histogram, cumulated for 8 different experiments, of the ratio of Eq. (2) left hand side (l.h.s.), 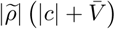 to right hand side (r.h.s.), 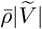. Red line: Gaussian fit, of average 1.11 (vertical dashes). *(B)* Histogram of the difference *ϕ_ρ_* – *ϕ_V_*. Red line: Gaussian fit, of average 3.12 (vertical dashes).

**Supporting Movie S1.**
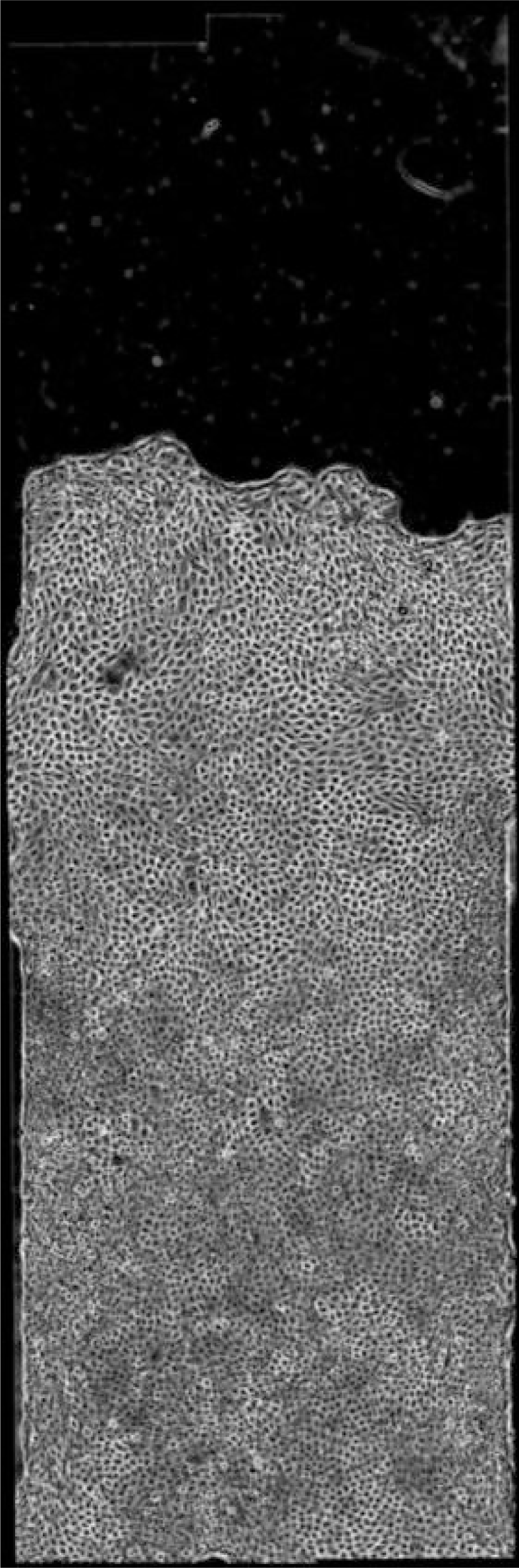
Monolayer of MDCK cells migrating within a straight strip with mitomycin C to prevent divisions. Phase-contrast image showing cell contours. Strip total length 4 mm, width 1 mm. The first image, noted *t* = 0, is taken after around 5 h of migration. Duration of the movie: 26 h. Because of file size constraints, the movie resolution has been decreased, and the time interval between frame has been doubled (10 min instead of 5 min in the original).

**Supporting Movie S2.**
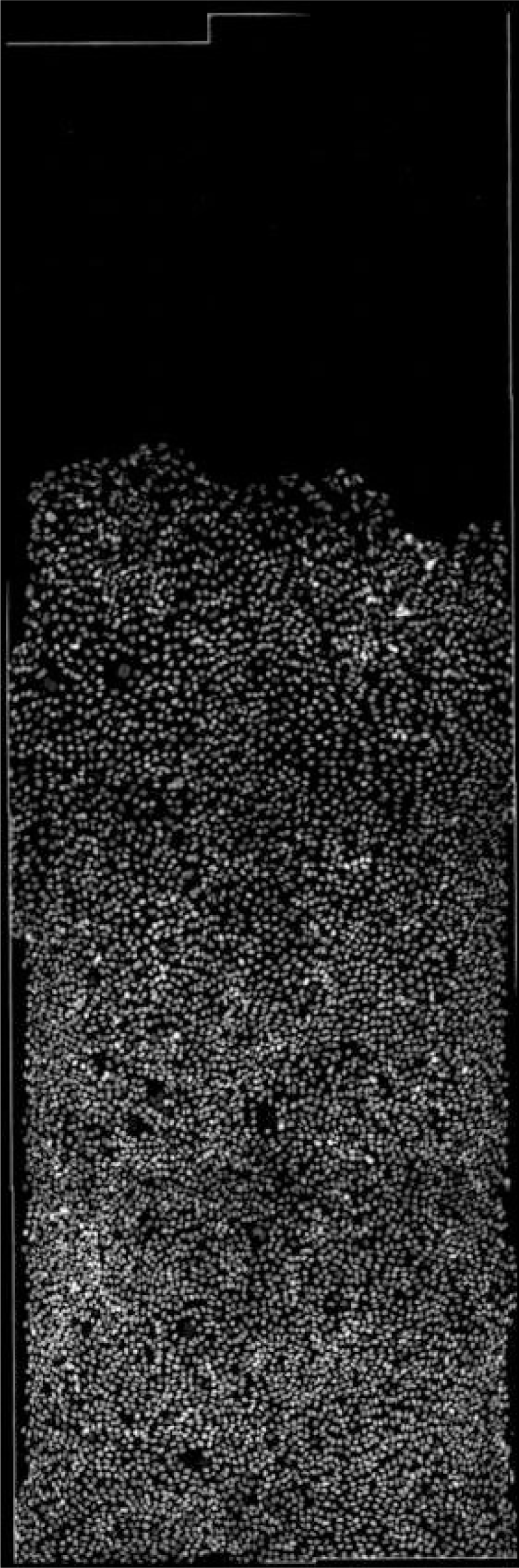
Same as Movie S1, nuclei labeled with histone GFP.

**Supporting Movie S3.**
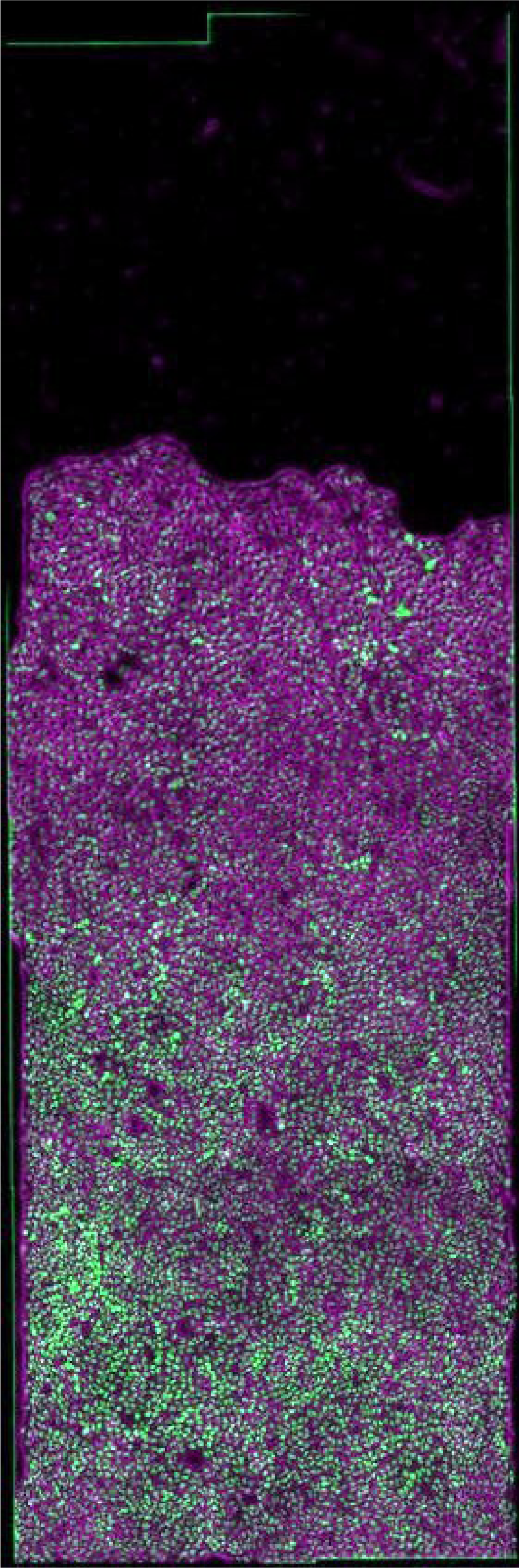
Merge of phase-contrast image showing cell contours (Movie S1) in purple, and nuclei labeled with histone GFP (Movie S2) in green.

**SI Supporting Movie S3.**
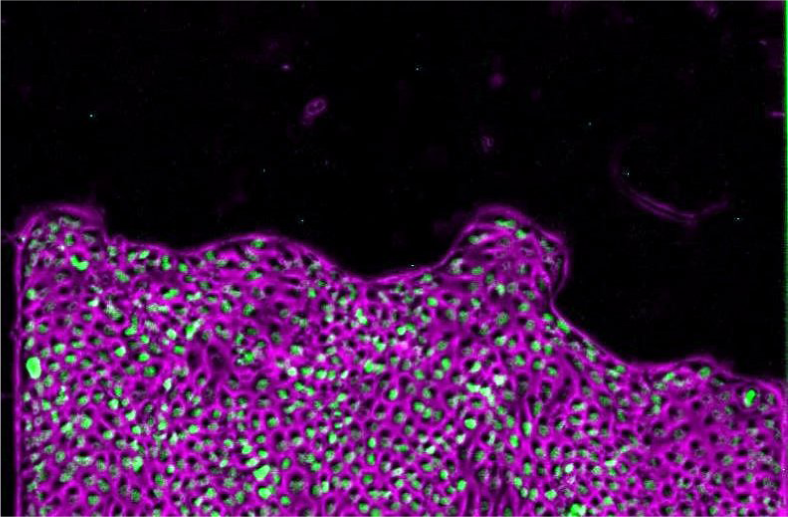
Same as Movie S3, with original time interval between frame (5 min), zoomed on the front.

**Supporting Movie S5.**
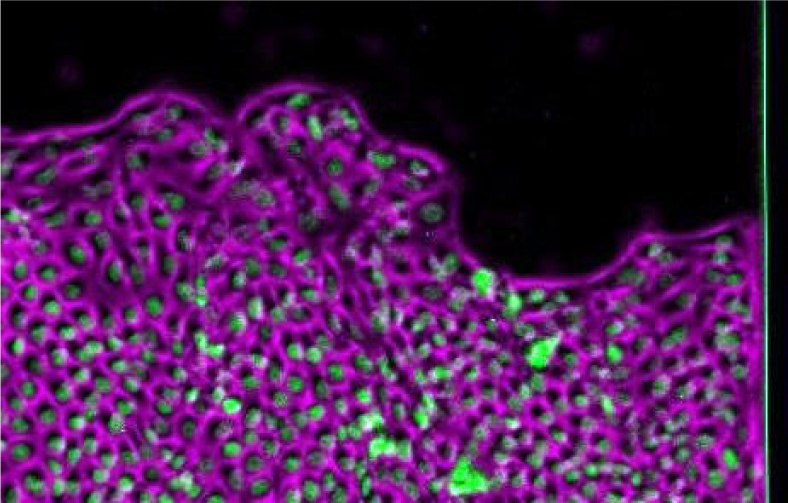
Same as Movie S3, with original time interval between frame (5 min), zoomed on the middle.

**Supporting Movie S6.**
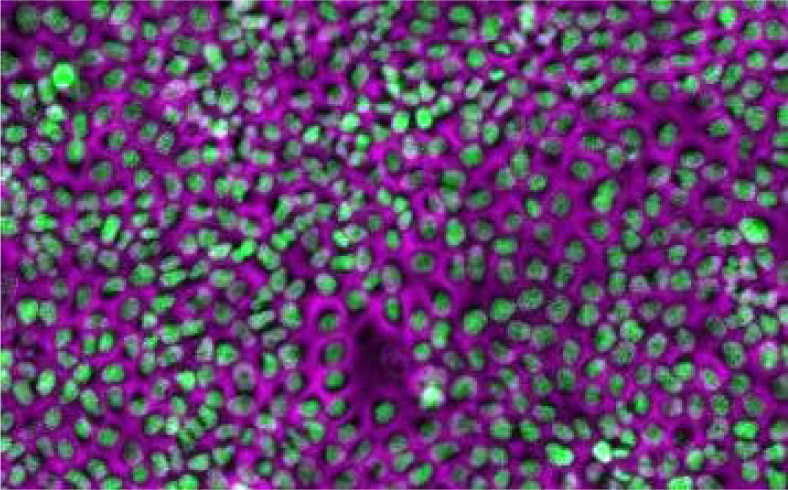
Same as Movie S3, with original time interval between frame (5 min), zoomed on the back.

**Supporting Movie S7.**
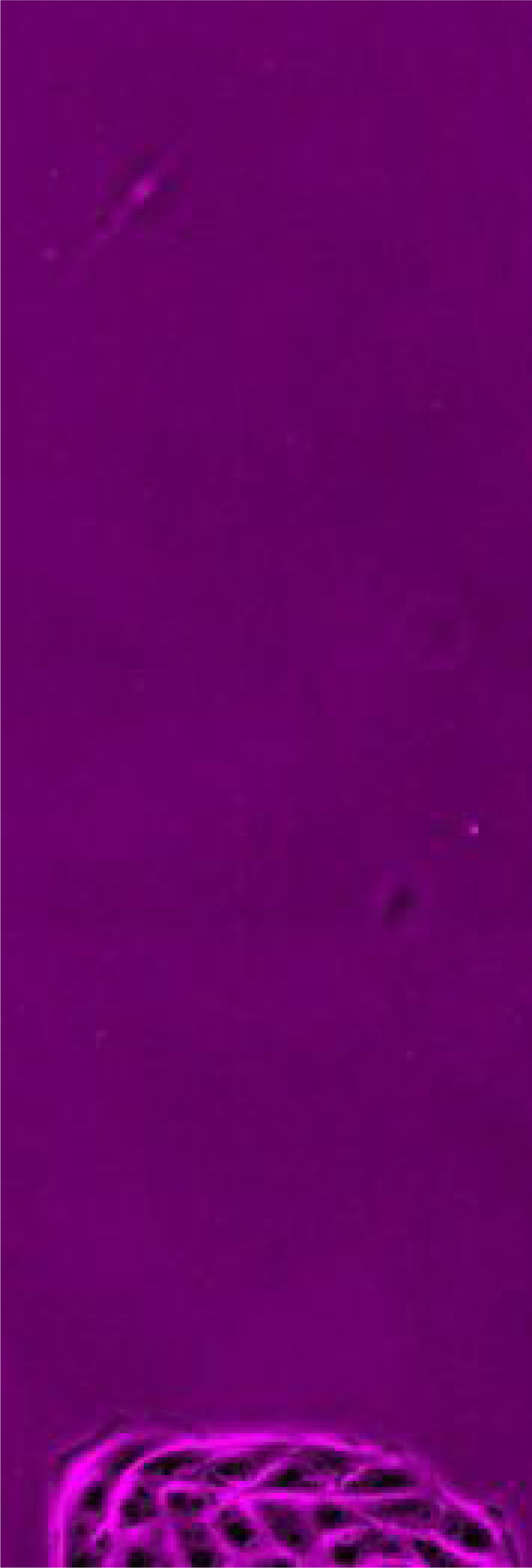
Same as Movie S3 (time interval between frames: 6 min). After four hours (corresponding to the missing image at the 10th second of the current movie), CK666 is added; lamellipodia (both cryptic and front ones) are no longer detectable.

